# The dynamic inositol phosphate network during development in *Drosophila melanogaster*

**DOI:** 10.64898/2026.06.18.733221

**Authors:** Anuj Shukla, Marvin Busch, Markus Häner, Anne-Kathrin Classen, Henning Jacob Jessen

## Abstract

Inositol phosphates (InsPs) and inositol pyrophosphates (PP-InsPs) are key regulators of cellular signaling and metabolism, yet developmental changes in the InsP/PP-InsP network in eukaryotic systems remain poorly understood. Here, we combine isomer-resolved capillary electrophoresis-mass spectrometry (CE-MS) with genetic perturbations to characterize InsP metabolism during the *Drosophila melanogaster* development. Our analyses reveal extensive developmental reprogramming of the InsP pathway, with early developmental stages exhibiting broad lower-order isomer diversity that progressively narrows into a more restricted metabolic state enriched in Ins(1,3,4,5,6)P_5_, InsP_6_, and PP-InsPs at later stages. Metamorphosis is characterized by a marked increase in 5-PP-InsP_5_, which emerges as the predominant pyrophosphate species in adults. Perturbation of dedicated kinases suggests that this metabolic remodeling is essential for developmental progression. RNAi-mediated IPK2 depletion disrupts higher-order InsP synthesis by reducing InsP_5_ levels, and results in failed metamorphosis. IPK1 depletion delays development and induces pronounced isomer-selective redistribution within the InsP_5_ pool, indicating network-level changes beyond simple precursor accumulation. IP6K depletion specifically abolishes 5-PP-InsP_5_ production, impairs tissue morphogenesis during pupal stages, and causes rapid post-eclosion lethality. Unexpectedly, depletion of either IPK2 or IP6K leads to the presence of 2-PP-InsP_5_. Collectively, these findings establish the *Drosophila melanogaster* InsP pathway as a dynamically regulated developmental network, particularly rich in previously undescribed InsP isomers, in which stage-specific metabolic remodeling parallels metamorphosis, tissue maturation, and adult viability.

## Introduction

Inositol phosphates (InsPs) and their pyrophosphorylated derivatives (PP-InsPs) are a class of highly phosphorylated metabolites derived from *myo*-inositol that function as versatile regulators of cellular communication, metabolism, and development across eukaryotes^1–3^. The six-carbon inositol ring provides a scaffold that can be differentially phosphorylated to encode several dozen predicted InsP isomers with distinct signaling functions^2^. Well-characterized members include Ins(1,4,5)P_3_, the calcium-release factor^4,5^, Ins(1,3,4,5,6)P_5_ produced from Ins(1,4,5)P_3_ through an inositol polyphosphate multikinase-dependent pathway implicated in tissue growth and viability^6^, InsP_6_, a ubiquitous storage and signaling molecule^7^, and the pyrophosphates 5-PP-InsP_5_ and 1,5-(PP)_2_-InsP_4_, which act as metabolic and signaling hubs in many organisms (**Fig. 1**)^8,9^.

**Figure 1.**
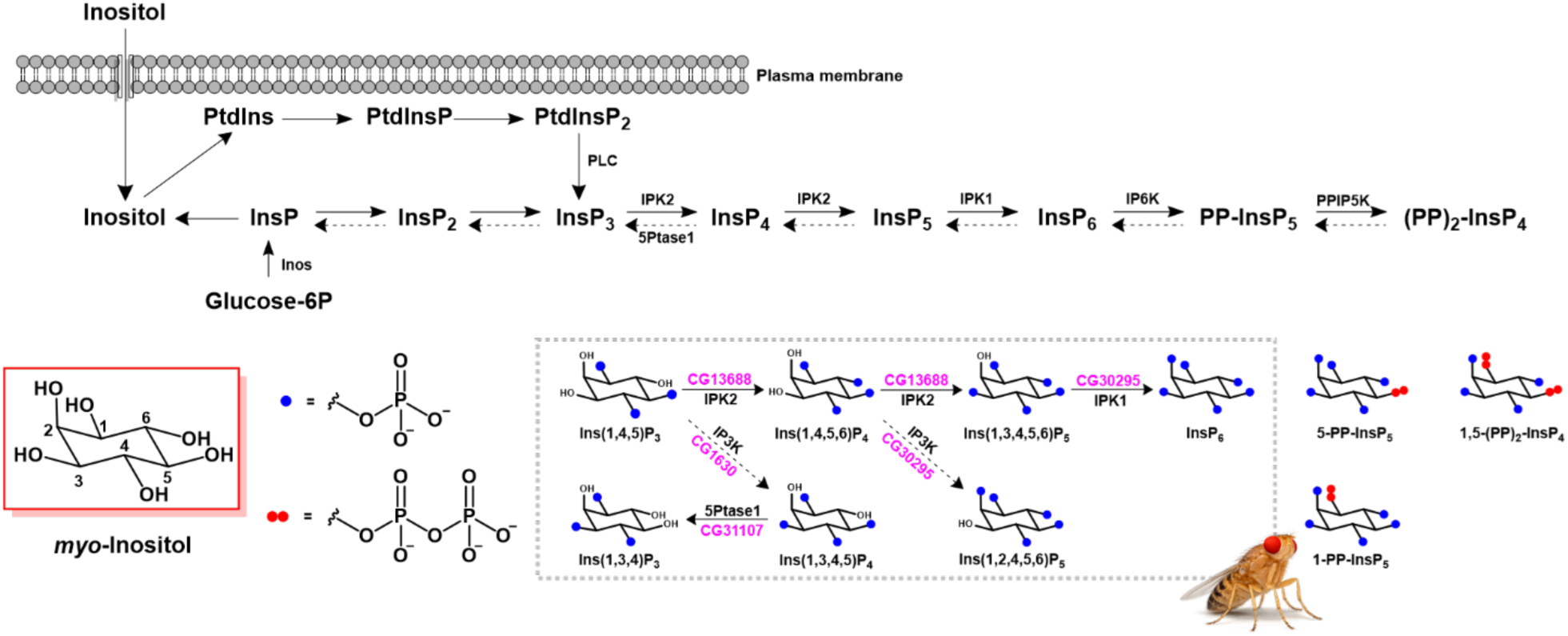
Conserved inositol phosphate and inositol pyrophosphate biosynthesis pathway with corresponding *Drosophila melanogaster* homologs. Phospholipase C (PLC) hydrolyzes PtdIns(4,5)P_2_ to Ins(1,4,5)P_3_, which is phosphorylated by inositol polyphosphate multikinase (IPMK/IPK2; CG13688) to generate InsP_4_ and InsP_5_ isomers. These intermediates are further phosphorylated by inositol pentakisphosphate 2-kinase (IPK1; CG30295) to produce InsP_6_. InsP_6_ is subsequently pyrophosphorylated by inositol hexakisphosphate kinase (IP6K) to form PP-InsP_5_, which can be further phosphorylated by diphosphoinositol pentakisphosphate kinase (PPIP5K) to generate (PP)_2_-InsP_4_. An alternative lipid-independent glucose-6-phosphate-dependent pathway, mediated by ITPK enzymes, can also contribute to the synthesis of higher InsPs. The scheme highlights conserved enzymatic steps and the corresponding *Drosophila* homologs in the dashed grey box. The red box shows the *myo*-inositol scaffold with carbon numbering used to define phosphorylation positions and isomer nomenclature.

The metabolic pathways leading to InsP production in different eukaryotic systems from yeasts to humans are broadly conserved (**Fig. 1**)^10^. In the canonical pathway, phospholipase C (PLC) hydrolyzes PtdIns(4,5)P_2_ to generate Ins(1,4,5)P_3_, which is phosphorylated at the 3 or 6 position by inositol polyphosphate multikinase (IPK2; also known as inositol polyphosphate multikinase, IPMK) to produce InsP_4_ isomers. These InsP_4_ species are further phosphorylated by IPK2 to form InsP_5_ isomers, followed by e.g. phosphorylation with inositol pentakisphosphate 2-kinase (IPK1) to generate InsP_6_. In addition to this lipid-dependent route, an alternative lipid-independent pathway proceeding through rearrangement of glucose-6-phosphate to Ins(3)P has been described, in which inositol-3-phosphate synthase (INO1/Inos) together with inositol trisphosphate kinases (ITPK) contribute to the generation of higher-order inositol phosphates (**Fig. 1**)^10^. Further, InsP_6_ – and to some extent InsP_5_ – serve as substrates for inositol hexakisphosphate kinases (IP6Ks), producing PP-InsP_5_ and PP-InsP_4_, which is subsequently phosphorylated by diphosphoinositol pentakisphosphate kinase (PP-IP5Ks) to generate (PP)_2_-InsP_4_ (**Fig. 1)**^10^. Its structure in mammals has been assigned as 1,5-(PP)_2_-InsP_4_^11^.

Since the discovery of InsP_3_ as a second messenger in animals in the 1980s^4^, InsPs have been implicated in diverse processes: nuclear mRNA export, phosphate sensing, and chromatin remodeling in yeast^12–15^; phosphate storage, stress responses, and hormone signaling in plants^16–19^ and proliferation, apoptosis, and insulin signaling in mammals^20–25^. Despite these major advances, little is known about InsP metabolism in invertebrate metazoans, a gap this study aims to address. *D. melanogaster*, with its well-known genetic toolkit and rapid development with a short life cycle, represents a promising system for dissecting InsP function in a multicellular developmental context. So far, only a few studies have used this organism to delineate the importance of the inositol phosphate network. Mutations in key pathway components including Plc21C, the InsP_3_ receptor, and IPK2 cause tissue-patterning defects and developmental arrest^6,26^. Notably, IPK2 has been shown to convert InsP_3_ to InsP_5_. The study links it to critical processes such as imaginal disc proliferation and survival via JAK/STAT signaling. Its loss causes pupal lethality^6^. Similarly, the enzyme Inos, encoding *myo*-inositol synthase, is important for proper morphological and metabolic development^27,28^. It may serve as a starting point for the lipid-independent pathway.

Despite these important findings, our understanding about general InsP metabolism in *D. melanogaster* remains very fragmented. This knowledge gap has been further spotlighted by a recent study describing a novel phosphate-sensing organelle: PXo bodies^29^. This organelle contains the fly homolog to human XPR1, a phosphate exporter^30^. Mammalian XPR1 has recently become the subject of diverse studies, demonstrating its structural rearrangement and regulation by the inositol pyrophosphate 1,5-(PP)_2_-InsP_4,_ which binds to its SPX-domains^31–33^. It is conceivable that PXo bodies are also regulated by PP-InsPs. Yet, information on InsP and in particular PP-InsP isomers and their turnover in *D. melanogaster* during its life cycle remain poorly characterized. Moreover, how changes in InsP concentrations relate to key developmental events such as larval growth, metamorphosis, and adult maturation remain unclear. Most previous studies have focused on genetic perturbations or measured only the total InsP levels, rather than identifying the specific positional isomers present at each developmental stage^6,34^. As a result, the lack of detailed isomer-level profiling has made it difficult to fully understand how InsP metabolic pathways are structured during development and how they influence stage-specific physiological processes.

One of the major problems in InsP research lies in the analytical challenges of probing and quantifying InsP metabolites and their interconversion^35^. InsPs occur at low concentrations, have multiple negative charges, lack intrinsic chromophores and exist as isobaric positional isomers, which complicates their separation, assignment, and quantification. Traditional methods such as SAX-HPLC with radiolabeling^36,37^ or PAGE^38,39^ with cationic-dye staining provide only partial resolution. Notably, radiolabeling approaches based on [^3^H]-inositol or [^32^P]-incorporation have enabled the detailed study of InsP metabolism particularly in plant, cultured cells, and *in vivo* fly systems^6,34^ by tracking the uptake and incorporation of labeled inositol^36,40–44^. However, these approaches have been technically challenging for positional isomer assignment and suffer from the potential downside of uneven labeling in whole organisms: they have therefore seen limited applications in *in vivo* animal studies.

Capillary electrophoresis electrospray ionization coupled to mass spectrometry (CE-ESI-MS) has been introduced as a selective method for the detection, assignment and sensitive quantitative analysis of InsPs and PP-InsPs^45–50^ . CE-ESI-MS allows charge-based separation of InsP isomers followed by high-resolution mass detection. This has made it possible to run quantitative and isomer-specific analyses of InsP_3_ to InsP_8_ from biological samples using heavy-isotope internal references^45,46,49,51^. It is a powerful tool for studying how InsP metabolism changes over time or under different physiological conditions and perturbations. Most notably, it can separate structurally similar isomers without radiolabeling and enables the delineation of metabolic fluxes^45–47,52^.

Because phosphorylation can occur at multiple positions around the inositol ring, several positional isomers and mirror image isomers (enantiomers) are possible (**Fig. 1**). The *myo*-inositol scaffold with carbon numbering (1–6) (red box in **Fig. 1**), defines phosphorylation positions and provides the basis for isomer nomenclature. CE-ESI-MS relies on an achiral separation and consequently enantiomers cannot be resolved (**Fig. S1**). For clarity in the figures, only one possible enantiomer is indicated without assignment of absolute stereochemistry. Accordingly, isomers are reported using lower-number nomenclature. For example, the enantiomeric pair Ins(1,4,5)P_3_/Ins(3,5,6)P_3_ is reported as Ins(1,4,5)P_3_ (**Fig. S1**).

In the following, we describe the use of CE-ESI-MS to systematically map different isomers of InsPs and PP-InsPs across egg, larval, pupal, and adult stages of *D. melanogaster*. By combining stage-specific profiling with RNAi-mediated knockdown of IPK2, IPK1 and IP6K, we uncover extensive developmental remodeling of the InsP network, identify some previously uncharacterized or invertebrate-enriched InsP and PP-InsP species, and reveal the crucial role of higher-order PP-InsPs during different developmental stages. Notably, depletion of 5-PP-InsP_5_ through IP6K knockdown results in defects during pupal to adult transition and reduced adult viability, indicating that proper regulation of PP-InsP metabolism is crucial during development.

Collectively, these findings provide the first isomer-resolved developmental network map of InsP isomers in an invertebrate model, giving a framework for comprehending the integration of dynamic remodeling of inositol phosphate metabolism with metabolic and developmental processes.

## Results

### 1. Developmental dynamics of inositol phosphate metabolism in *D. melanogaster*

*D. melanogasteŕs* life cycle is divided into discrete developmental stages that are marked by changes in cell architecture, metabolic demand and tissue organization. Embryogenesis establishes the basic body structure, larval stages are dominated by rapid growth and nutrient accumulation, and pupation involves extensive metamorphic remodeling, during which larval tissues are eliminated and adult structures are formed^53^. Further, the emerging adults undergo a period of post-eclosion maturation before reaching physiological homeostasis.

To determine how InsP/PP-InsP metabolism is regulated during these development transitions, we performed specific analyses across eight key stages of the wild-type *D. melanogaster* life cycle (w^1118^ background): 1) eggs; 2-4) first-, second-, and third-instar larvae; 5-6) early and late pupae; 7) newly eclosed adults (3-5 days); 8) mature adults (15-20 days). Tissue composition, hydration and protein content vary substantially across development (**Supplementary page 2**). For consistency, the quantitative comparisons of InsP metabolites were performed using fresh weight normalization across development stages^54^. InsPs and PP-InsPs were obtained using perchloric acid-based extraction followed by TiO_2_-mediated enrichment, a strategy shown to provide robust recovery of highly phosphorylated metabolites, including InsPs and PP-InsPs, from complex biological samples^49,55^.

During the analyses, we encountered an unexpectedly large array of InsPs and PP-InsPs. Therefore, we first studied, whether *myo*-inositol is the predominant form found in *D. melanogaster* or whether there are also significant amounts of *scyllo*-inositol present. CE-ESI-MS analysis of these two isomers (**Fig. S2**) confirmed that more than 95% of inositol released from inositol phosphates by phytase or heat treatment has likely *myo*-inositol configuration and that less than 5% are derived from *scyllo*-inositol, consistent with canonical eukaryotic InsP metabolism^56^. Other potential isomers were not investigated due to their low natural abundance. We also evaluated extraction recovery for different isomers across development stages, which demonstrated an excellent recovery of 75-80% (**Fig. S3**). ATP levels were quantified in parallel across developmental stages and normalized to fresh tissue weight (**Fig. S4**).

With this information, we profiled *myo*-InsP isomers using high-resolution CE-ESI-MS. Analysis of whole-organism extracts revealed pronounced developmental dynamics, with early stages dominated by lower-order InsPs and later stages characterized by progressive enrichment of InsP_6_ and the emergence of PP-InsPs (**Fig. 2** and **Fig. S5-12**). Up to twenty different isomers of InsPs and PP-InsPs were observed. This is in contrast to previous studies where only up to nine InsPs species, were reported^6,34^. These findings indicate that the complexity of the *Drosophila melanogaster* InsP network has so far been underestimated.

**Figure 2.**
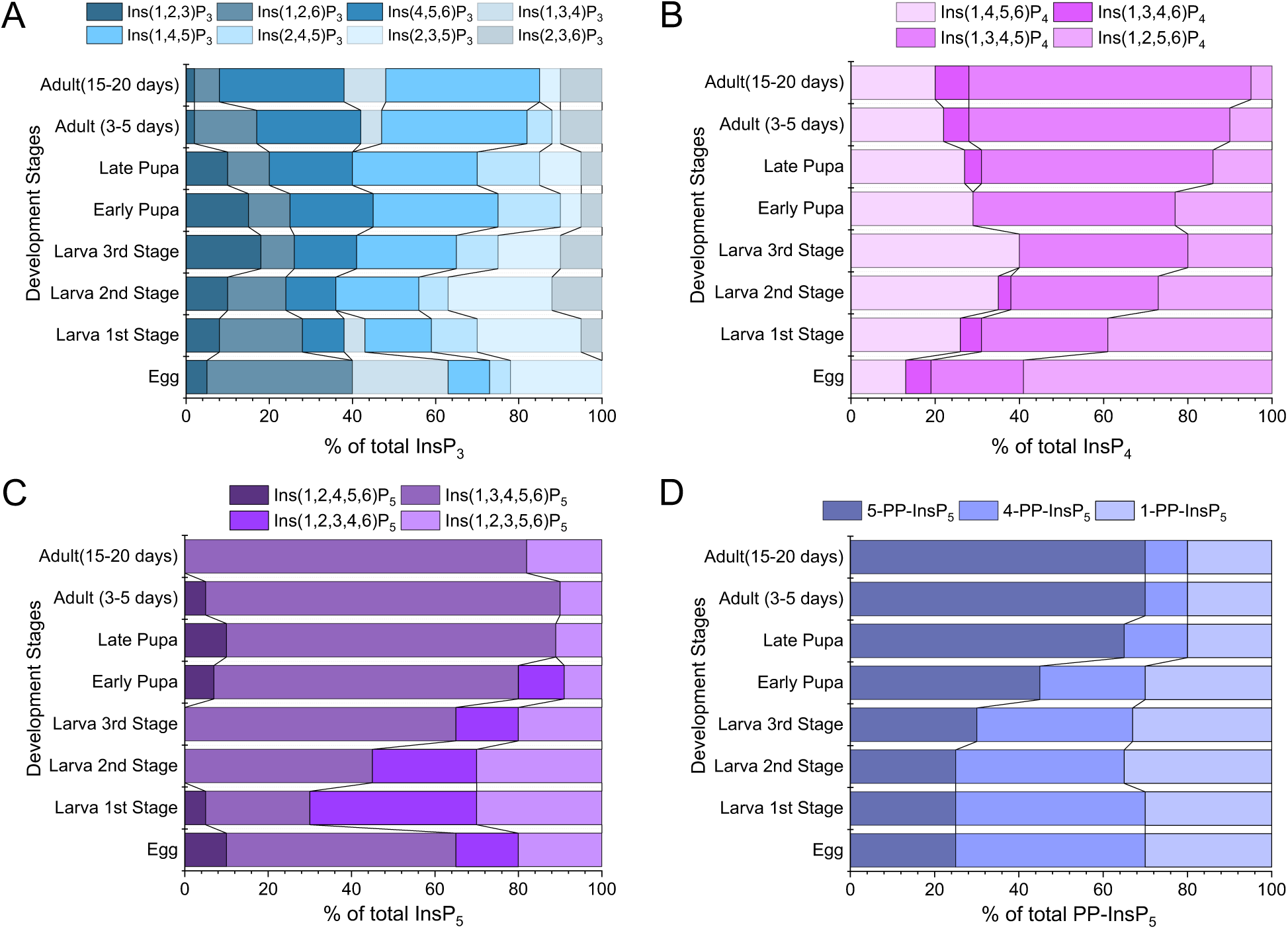
Stage-specific remodeling of inositol phosphate metabolism during *D. melanogaster* development. Stacked bar plots showing the relative composition (percentage of total signal within each InsP class) of individual isomers of (**A**) InsP_3_, (**B**) InsP_4_, (**C**) InsP_5_, and (**D**) PP-InsP_5_ across developmental stages (egg; 1^st^,2^nd^,3^rd^ instar larvae; early and late pupae; 3–5 days adults; and mature adults (15–20 days). Early developmental stages are enriched in Ins(1,2,6)P_3_ and Ins(2,3,5)P_3_, whereas the transition to late larval and pupal stages is characterized by increased Ins(1,4,5)P_3_ and Ins(4,5,6)P_3_. The InsP_4_ pool shifts from Ins(1,2,5,6)P_4_ in early stages towards Ins(1,3,4,5)P_4_ and Ins(1,4,5,6)P_4_ during larval-pupal development, with Ins(1,3,4,5)P_4_ becoming the prominent species in adults. Among InsP₅ species, Ins(1,3,4,5,6)P_5_ increases strongly from the third-instar larva to adult, while Ins(1,2,3,5,6)P_5_/Ins(1,2,3,4,5)P_5_ remains minor and Ins(1,2,3,4,6)P_5_ is detectable in very low amounts only. Within the pyrophosphate pool, 1-PP-InsP_5_ remains relatively stable during development. In contrast, 5-PP-InsP_5_ increases sharply-while 4-PP-InsP_5_ declines- from late larvae stages through adult flies, and results in a pronounced enrichment of inositol pyrophosphates in adult flies. Values were normalized to stable isotopic internal standards (^13^C_6_/^18^O_12_) and tissue mass. Data are shown as mean values from n = 3 independent biological replicates per developmental stage and standard deviations are reported in **Table S2**. Only one possible enantiomer is indicated for clarity.

#### 1.1 Dynamic remodeling of InsP_3_ to InsP_6_ during development

To enable confident positional isomer assignment across InsP_3_-InsP_5,_ we employed a two-step reference strategy. Where available, InsP_3_-InsP_5_ isomers were assigned by co-migration with stable isotope-labeled [^13^C_6_] standards^46,47^. These [^13^C_6_]-labeled standards were additionally used to assign isomer-mixtures of InsP_3_-InsP_5_ obtained by pyrohydrolysis of [^18^O_12_]-InsP_6_. These mixtures, used as internal references, were assigned by co-migration with the corresponding [^13^C_6_] standards and commercial InsPs (**Fig. S5A-C**). Due to the diversity of detected InsP_3_ isomers in samples, we also synthesized a new ^18^O_6_-labeled reference compound to assign Ins(2,4,6)P_3_ (**Fig. S5D**). Co-migration experiments confirmed its separation from other InsP_3_ isomers under our CE-ESI-MS conditions, enabling assignment (**Fig. S5E**).

Ins(1,4,5)P_3_ is one of the most extensively studied species in the inositol phosphate pathway, known as the calcium-release factor in animal cells^57–59^. In wild-type (WT), eight different positional InsP_3_ isomers were detected during CE-ESI-MS analysis that were assigned through comparison with products of a pyrohydrolysed ^18^O_12_-InsP_6_ standard (**Fig. S7A-D**). To our knowledge, this represents the largest number of InsP_3_ isomers reported in a single organismal system to date^42^. Quantitative profiling revealed pronounced, developmental remodeling of the InsP_3_ pool (**Fig. 2A**). In eggs, Ins(1,2,6)P_3_ dominated, accounting for more than one-third of total InsP_3_, with additional contributions from Ins(1,3,4)P_3_ (∼20%), Ins(2,3,5)P_3_ (∼20%) and Ins(1,4,5)P_3_ (∼10%). The low abundance isomers Ins(2,4,5)P_3_ and Ins(1,2,3)P_3_ were also detected. During larval development, a sharp decrease in the abundance of Ins(1,2,6)P_3_ and Ins(2,3,5)P_3_ was detected, accompanied by a prominent increase in Ins(1,4,5)P_3_, Ins(4,5,6)P_3_, and Ins(1,2,3)P_3_. A second transition occurred during pupation, when Ins(1,4,5)P_3_ emerged as the dominant InsP_3_ species, reaching approximately 40% of the total InsP_3_ pool in the late pupa. This was paralleled by an enrichment of Ins(4,5,6)P_3_ (∼20%) and Ins(2,4,5)P_3_ (∼15%). In contrast, isomers formerly enriched in larvae were strongly reduced in adults and the InsP_3_ profile stabilized into a reproducible composition characterized by dominance of Ins(1,4,5)P_3_ and Ins(4,5,6)P_3_; these species accounted for more than 70% of the total InsP_3_ pool (**Fig. 2A**).

Next, we scrutinized inositol tetrakisphosphates (InsP_4_), which function both as intermediates in higher-order InsP biosynthesis and as regulators of diverse cellular processes or cofactors^3,24,60–65^. CE-ESI-MS analysis resolved four positional isomers across different development stages: Ins(1,4,5,6)P_4_, Ins(1,3,4,6)P_4_, Ins(1,3,4,5)P_4_, and Ins(1,2,5,6)P_4_, of which only Ins(1,4,5,6)P_4_ and Ins(1,3,4,5)P_4_ have been reported previously in *D. melanogaster*^34^ (**Fig. S8A-D**). Quantitative profiling revealed pronounced, stage-specific remodeling of the InsP_4_ pool during *D. melanogaster* development. In eggs, the InsP_4_ pool was dominated by Ins(1,2,5,6)P_4_, which accounted for ∼ 60% of total InsP_4_, with additional contributions from Ins(1,3,4,5)P_4_ (∼20%) and Ins(1,4,5,6)P_4_ (∼13%). During larval development, this profile underwent a progressive redistribution, characterized by a steady decline in Ins(1,2,5,6)P_4_ and a prominent enrichment in Ins(1,3,4,5)P_4_ and Ins(1,4,5,6)P_4_, which contributed ca. 80% of the total InsP_4_ pool by the third instar stage. During pupation and the adult phase, a second remodeling occurred, in which Ins(1,2,4,5)P_4_ levels gradually declined to ∼10% and Ins(1,3,4,5)P_4_ emerged as a major component contributing more than 60% of the total InsP_4_ pool (**Fig. 2B**).

We subsequently examined inositol pentakisphosphate (InsP_5_) isomers during development. Ins(1,3,4,5,6)P_5_ serves as a major biosynthetic intermediate in eukaryotic systems^3,24,66^. The Ins(1,3,4,5,6)P_5_ isomer was identified and quantified by co-migration with a ^13^C_6_-labeled internal standard^49,67^ (**Fig. S9A-D**), while the other Ins(1,2,4,5,6)P_5_, Ins(1,2,3,4,6)P_5_ and Ins(1,2,3,4,5)P_5_ species were confirmed by comparison with products generated from pyrohydrolysis of an ^18^O_12_-InsP_6_ standard^68,69^ (**Fig. S9A**). Quantitative profiling revealed pronounced developmental changes in the InsP_5_ pool. In eggs, Ins(1,3,4,5,6)P_5_ accounted for just over half of the total InsP_5_ pool with substantial contributions from Ins(1,2,3,4,6)P_5_ (∼25%), and Ins(1,2,3,4,5)P_5_ (∼20%). During early larval development, a transient reduction in Ins(1,3,4,5,6)P_5_ and a corresponding increase in other InsP_5_ isomers was observed, which together amounted to ∼75% of the total InsP_5_ pool. However, from the second instar we observed an enrichment in the Ins(1,3,4,5,6)P_5_ levels, which reached around 65% by third instar and ∼80-95% in pupa and adults (**Fig. 2C**).

Inositol hexakisphosphate (InsP_6_) is the fully phosphorylated form of inositol and a central precursor for the inositol pyrophosphates^1,2^. InsP_6_ was identified and quantified by co-migration with a ^13^C_6_-labeled internal standard^49,67^ (**Fig. S10A**). InsP_6_ abundance was quantified directly without isomeric deconvolution and is reported in absolute amounts normalized to fresh weight (pmol mg⁻¹ FW) in the supplementary information. In our analysis, it showed stage-specific variation during development (**Fig. S10B**) with several-fold changes between stages.

#### 1.2 Enrichment of inositol pyrophosphates during pupal transitions

We next examined inositol pyrophosphates (PP-InsPs), which act as second messengers and regulators of phosphate and energy homeostasis in diverse organisms^2,70^. CE-MS resolved three PP-InsP_5_ isomers: 5-PP-InsP_5_, 1-PP-InsP_5_, and 4-PP-InsP_5_ along with a single detectable (PP)_2_-InsP_4_: 1,5-(PP)_2_-InsP_4_. Their identities and separation were confirmed by co-migration with authentic ^13^C_6_-labeled 5-PP-InsP_5_ and 1-PP-InsP_5_, ^18^O_2_-labeled 4-PP-InsP_5_ (Fig S6A, S11A-D)^45,68,71^, and ^13^C_6_ 1,5-(PP)_2_-InsP_4_^49,67^. As baseline separation of isobaric 5-PP-InsP_5_ and 4-PP-InsP_5_ was not fully achieved under all developmental conditions based on matrix effects (**Fig. S11A-D**), individual reported isomer abundances should be considered as estimates; nevertheless, consistent stage-dependent shifts in peak areas were reproducibly observed across biological replicates. Further, to exclude potential isomerization during sample extraction, preparation or analysis, pre-spiking experiments were performed using ^13^C_6_-labeled 5-PP-InsP_5_ and 1-PP-InsP_5_ standards, confirming that no additional PP-InsP isomers were produced under the extraction conditions (**Fig. S6B**).

During early development (eggs, larvae), the PP-InsP_5_ pool was dominated by 4-PP-InsP_5_ (40-45%) and 1-PP-InsP_5_ (30-35%) (**Fig. 2D**). In contrast to this early PP-InsP distribution, 5-PP-InsP_5_ increased markedly from the late larvae and early pupae stages and becomes the dominant species (∼70%) in adults (15-20 days), accompanied by a strong decline in 4-PP-InsP_5_. Our analysis highlights the prominent accumulation of 4-PP-InsP_5_ in *Drosophila melanogaster*, a PP-InsP isomer that is currently detectable only by CE-MS. In addition, a prominent single (PP)_2_-InsP_4_, assigned as 1,5-(PP)_2_-InsP_4_ by co-migration with a heavy isotope reference, was detected at low absolute amounts of ca. 0.1 pmol mg⁻¹ FW in adults - approximately two orders of magnitude lower than the InsP_6_ levels (**Fig. S10B and S12B**). During development, we have also detected the 4,5-(PP)_2_-InsP_4_; however, its abundance remained close to the detection limit and was therefore not quantitatively evaluated. In contrast, 1,5-(PP)_2_-InsP_4_ followed a pattern similar to that of 5-PP-InsP_5_ with a marked increase in early pupa and then remaining relatively constant throughout later development (**Fig. S12B**).

### 2. InsP kinases orchestrate developmental progression through distinct metabolic roles

#### 2.1 IPK2 knockdown disrupts InsP flux and causes larval arrest

We sought to determine how RNAi-mediated depletion of IPK2 affects the inositol phosphate network during development and performed systemic RNAi knockdowns under the control of a ubiquitous tub-GAL4 driver. For our analyses, we examined InsP composition by CE-MS in pupae at 12-24 h after puparium formation, a developmental window in which our wild-type profiling had revealed pronounced changes in InsP composition. Consistent with the severe developmental phenotypes, 72% of IPK2 RNAi pupae failed to eclose. As our focus was on the InsP network, we did not further characterize morphological phenotypes. Strikingly, RNAi-mediated depletion of IPK2 in pupae revealed extensive changes within the whole InsP network, with pronounced alterations in InsP_3_, InsP_4_, InsP_5_, and PP-InsP levels (**Fig. 3, S14A-C**).

**Figure 3.**
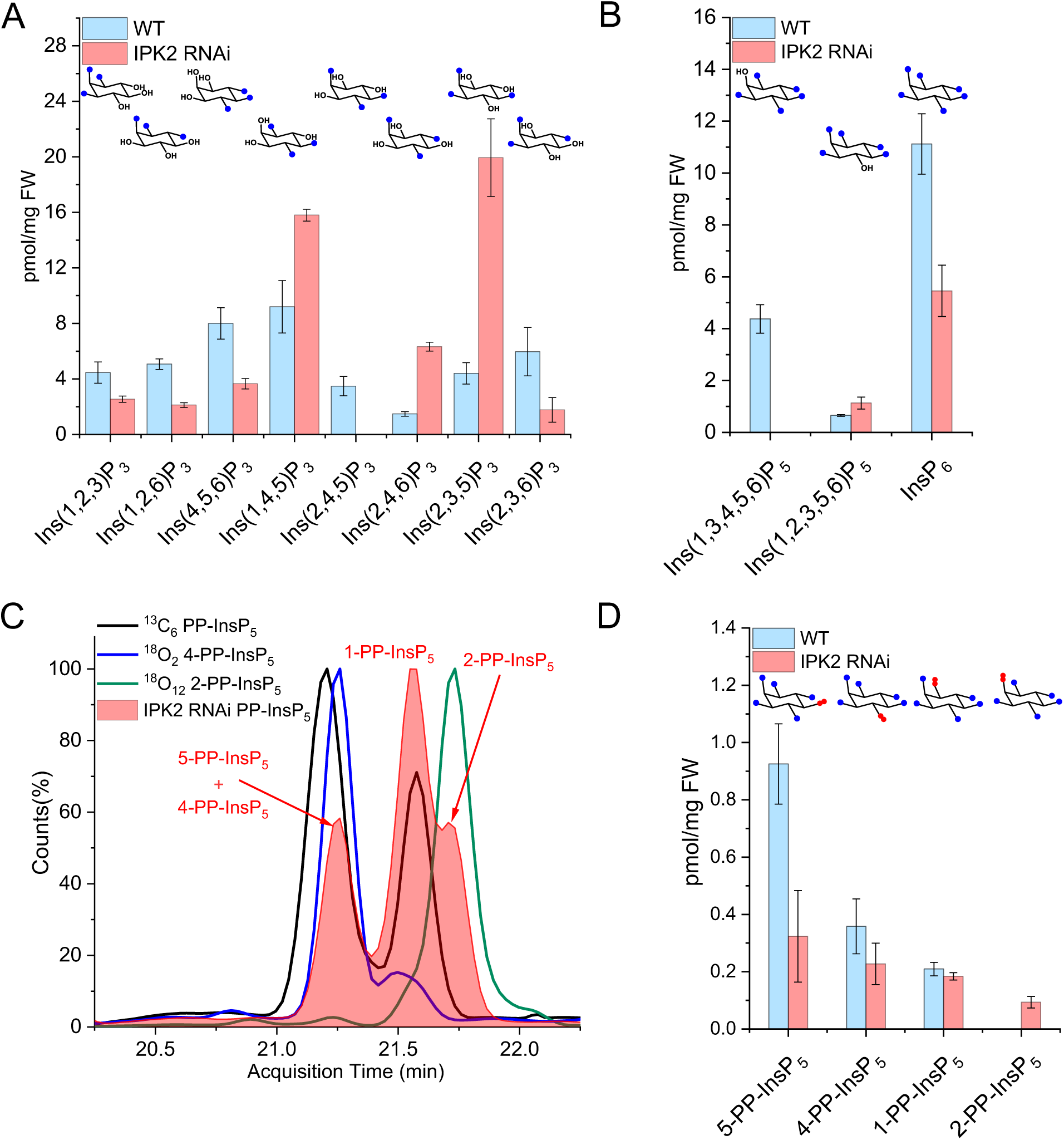
Changes in inositol phosphate abundance and identity upon IPK2 depletion in *D. melanogaster* pupae. **(A)** Quantification of InsP_3_ isomers in 12-24 h pupae following RNAi-mediated depletion of IPK2, measured by CE-ESI-MS after perchloric acid extraction and TiO_2_ enrichment. Bars represent mean ± s.d. (n=3). Wild-type samples are shown in sky blue, whereas IPK2 RNAi samples are shown in red, illustrating changes in the InsP_3_ isomer composition upon IPK2 depletion. **(B)** Quantification of InsP_5_ and InsP_6_ levels in the same samples. Bars represent mean ± s.d. (n=3) (WT in sky blue; IPK2 RNAi in red) and show the relative abundance of higher inositol phosphates following IPK2 depletion. **(C)** Representative CE-MS electropherogram showing separation of PP-InsP_5_ isomers. The red filled trace corresponds to endogenous PP-InsP_5_ signals detected in IPK2-depleted samples, whereas the black line represents [^13^C_6_]-labeled reference standards for 5-PP-InsP_5_ (migration time 21.3 min) and 1-PP-InsP_5_ (migration time 21.6 min). The blue line corresponds to the ^18^O_2_ 4-PP-InsP_5_ reference and the green line represents the ^18^O_12_ 2-PP-InsP_5_ reference. Enhanced separation of 5-PP-InsP_5_ and 4-PP-InsP_5_ is shown in **Fig. S14D**. **(D)** Quantification of PP-InsP_5_ isomers in 12–24 h pupae following IPK2 depletion. Bars represent mean ± s.d. (n=3) (WT in sky blue; IPK2 RNAi in red).

In WT pupae, Ins(1,4,5)P_3_ (ca. 9 pmol/mg FW) is one of the most abundant InsP_3_ isomers, together with Ins(1,2,6)P_3_ and Ins(2,3,5)P_3_ (**Fig. 3A & S13A**). Upon RNAi-mediated depletion of IPK2 in pupae, the total InsP_3_ pool was increased by approximately 20%. Ins(1,4,5)P_3_ increased by 2-fold, Ins(2,4,6)P_3_ by 3-fold and Ins(2,3,5)P_3_ by more than 6-fold. In contrast, Ins(2,4,5)P_3_ was absent and the levels of most other assigned InsP_3_ isomers were markedly reduced (**Fig 3A, S14A**). At the InsP_4_ level, Ins(1,3,4,5)P_4_ was markedly reduced, while Ins(1,2,5,6)P_4_ accumulated significantly (**Fig S13B, S14B**). Accurate assignment of several new InsP_4_ isomers was not possible using the pyrohydrolysis reference mixture^47^, as many of the possible isomers are not yet available in pure form. Nevertheless, it becomes apparent that RNAi-mediated depletion of IPK2 results in extensive remodeling of the InsP_3_ and InsP_4_ pool, consistent with its broad substrate specificity.

Furthermore, InsP_5_ and InsP_6_ were also markedly affected by IPK2 depletion. In wild-type pupae, CE-MS resolves two InsP_5_ isomers, including Ins(1,3,4,5,6)P_5_, and Ins(1,2,3,5,6)P_5_, and InsP_6_ (**Fig. 3B, S13C**). In the IPK2 RNAi, a 60-70% reduction in total InsP_6_ and InsP_5_ levels is observed (**Fig. 3B**). Notably, Ins(1,3,4,5,6)P_5_, the canonical precursor to InsP_6_, and product of InsP_4_ to InsP_5_ conversion by IPK2, was nearly undetectable and the residual InsP_5_ pool was almost exclusively composed of Ins(1,2,3,5,6)P_5_ (**Fig. S14C**). This redistribution was accompanied by an approximately two-fold reduction in InsP_6_ levels (**Fig. 3B**), indicating impaired synthesis of the fully phosphorylated state from the available isomers.

We also observed significant effects on PP-InsPs. Wild-type pupae and adults consistently contained 5-PP-InsP_5_, 1-PP-InsP_5_, and 4-PP-InsP_5_ (**Fig. S13D**). Upon RNAi-mediated depletion of IPK2 in pupae, loss of 5-PP-InsP_5_ (ca. 50-60% decrease) was observed (**Fig. 3D, S14D**), paralleled by reduction of 4-PP-InsP_5_, yet to a lower extent (**Fig. 3D**). 1-PP-InsP_5_ was not affected (**Fig. 3C-D, S14D**). Notably, an additional, PP-InsP_5_ peak emerged in IPK2-depleted pupae that was absent in the control. Based on prior reports^45^ describing the existence of non-canonical PP-InsP_5_ isomers, 2-PP-InsP_5_ was considered a candidate. To test this, we synthesized an [^18^O_12_] 2-PP-InsP_5_ reference^51^ (**Fig. S15A-B**) and performed post-spiking experiments, enabling comparison of migration time with the endogenous signal (**Fig. 3C**). The endogenous peak co-migrated precisely with the ^18^O-labeled reference, and displayed the accurate mass and fragmentation pattern for a PP-InsP_5_, enabling assignment of this species as 2-PP-InsP_5_ (**Fig. 3C-D**). The origin of this isomer in the IPK2-depleted background remains unclear.

#### 2.2 IPK1-depletion affects the inositol phosphate network

Inositol pentakisphosphate 2-kinase (IPK1) phosphorylates Ins(1,3,4,5,6)P_5_ to InsP_6_ in *D. melanogaster* (**Fig. 1**)^34^. RNAi-mediated depletion of IPK1 resulted in a reproducible 12-24 h delay in pupal formation compared to wild-type and subsequent pupal to adult transition was also delayed (**Fig. S14G-H**).

Metabolic profiling of RNAi-mediated IPK1-depleted pupae by CE-MS revealed specific changes in the inositol phosphate network. In wild-type pupae, Ins(1,4,5)P_3_ (ca. 9 pmol/mg FW) occurred as one of the most abundant InsP_3_ isomers, together with Ins(1,2,6)P_3_ and Ins(2,3,5)P_3_ (**Fig. 4A & S13A**). RNAi-mediated depletion of IPK1 in pupae results in a redistribution of the InsP_3_ isomers without significantly altering the total concentration. Notably, Ins(4,5,6)P_3_ accumulated 3-fold compared to wild-type whereas almost no change was observed in Ins(1,4,5)P_3_. Ins(1,2,6)P_3_ decreased 2-fold and several other InsP_3_ species, such as Ins(1,2,3)P_3_, Ins(2,3,5)P_3_, and Ins(2,3,6)P_3_, were strongly diminished (**Fig. 4A & S14E**). At the InsP_4_ level, multiple isomers were resolved, with reduced Ins(1,2,4,5)P_4_ and Ins(1,3,4,5)P_4_ (**Fig. S14F**) relative to wild-type pupae (**Fig. S13B**). This was accompanied by accumulation of Ins(1,3,4,6)P_4_ and Ins(1,4,5,6)P_4_ and additional unassigned isomers (**Fig. S14F**). Several peaks could not be assigned due to limited availability of reference compounds (**Fig. S13B**).

**Figure 4.**
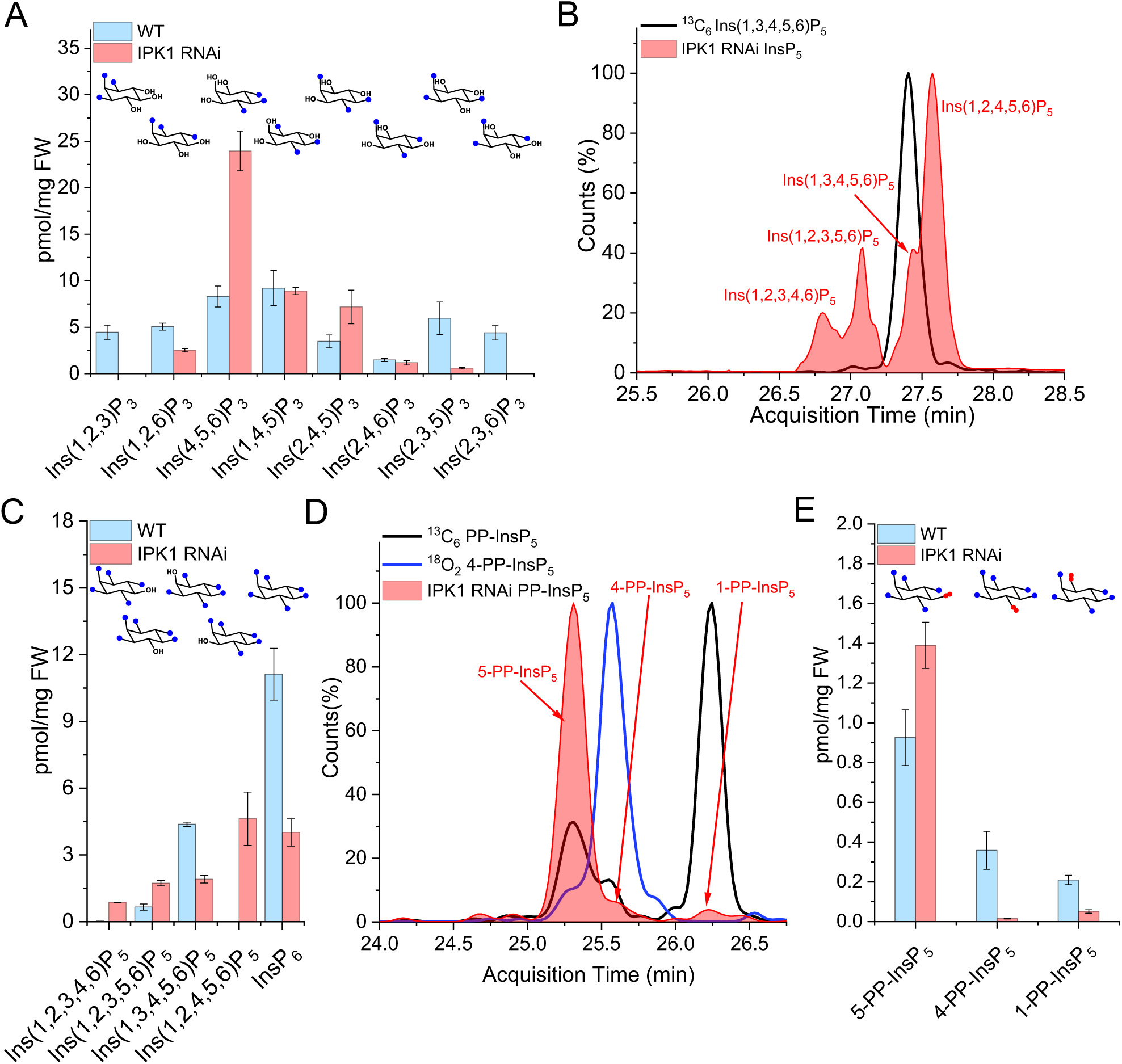
Remodeling of inositol phosphate metabolism upon IPK1 depletion in *D. melanogaster* pupae. (A) Quantification of InsP_3_ isomers in 12–24 h pupae following RNAi-mediated depletion of IPK1, measured by CE-MS after perchloric acid extraction and TiO_2_ enrichment. Bars represent mean ± s.d. (n=3). Wild-type samples are shown in sky blue, whereas IPK1 RNAi samples are shown in red. (B) Representative CE-MS extracted ion electropherogram showing separation of InsP_5_ positional isomers detected in IPK1-depleted pupa (red). Individual peaks correspond to Ins(1,2,3,4,6)P_5_, Ins(1,2,3,5,6)P_5_, Ins(1,3,4,5,6)P_5_, and Ins(1,2,4,5,6)P_5_. (C) Quantification of InsP_5_ isomers and InsP_6_ in 12–24 h pupae following IPK1 depletion. Bars represent mean ± s.d. (n=3) (WT in sky blue; IPK1 RNAi in red), showing accumulation of specific InsP₅ species accompanied by reduced InsP_6_ levels upon IPK1 depletion. (D) Representative CE-MS electropherogram showing separation of PP-InsP_5_ isomers. The red filled trace corresponds to endogenous PP-InsP_5_ signals detected in pupal extracts, whereas the blue line represents the ^18^O-labeled 4-PP-InsP_5_ reference and the black line corresponds to [^13^C_6_]-labeled 5-PP-InsP_5_ (migration time 25.3 min) and 1-PP-InsP_5_ (migration time 26.3 min) standards. (E) Quantification of PP-InsP_5_ isomers in 12–24 h pupae following IPK1 depletion. Bars represent mean ± s.d. (n=3) (WT in sky blue; IPK1 RNAi in red).

IPK1 depletion in pupae also had a pronounced effect on the InsP_5_ and InsP_6_ pools (**Fig. 4B-C**). In wild-type pupae, the InsP_5_ pool was dominated by Ins(1,3,4,5,6)P_5_ (ca. 4.5 pmol/mg FW) and also contained Ins(1,2,3,5,6)P_5_ (ca. 0.8 pmol/mg FW). RNAi-mediated IPK1 depletion in pupae resulted in a 2-fold reduction of Ins(1,3,4,5,6)P_5_ (ca. 1.9 pmol/mg FW), while Ins(1,2,4,5,6)P_5_ accumulated markedly to ca. 4.6 pmol/mg FW. In addition, Ins(1,2,3,4,6)P_5_ levels increased ca. 2-fold. Overall, the total InsP_5_ pool doubled upon IPK1 depletion as compared to wild-type controls (**Fig. 4 B-C and S13C**). In line with expectations regarding the role of IPK1, the InsP_6_ level was reduced by ca. 50-60% relative to wild-type (**Fig 4C**). Given the still significant abundance of InsP_6_, other kinases might complement its generation. Another key change was the ca. 80-90% reduction in 1-PP-InsP_5_ and 4-PP-InsP_5_ levels that was accompanied by a marked increase in 5-PP-InsP_5_ levels (**Fig. 4D-E**).

#### 2.3 IP6K-dependent 5-PP-InsP_5_ is essential for tissue remodeling during pupal stages and adult viability

While IP6K enzymes are well established as the primary enzymes to generate 5-PP-InsP_5_ in yeast and mammals, their function in *D. melanogaster* has not yet been experimentally studied. An early genetic and biochemical analysis of the *D. melanogaster* inositol phosphate pathway^34^ identified CG10082 as a putative IP6K homolog based on sequence similarity to mammalian IP6-kinases (**Fig. 1 & S16**)^34^. To investigate the functional role of IP6K *in vivo*, we performed systemic RNAi-mediated depletion and examined inositol pyrophosphate levels in pupae. In wild-type, 5-PP-InsP_5_ levels normally increase during this stage and become the most abundant PP-InsP_5_ based on our profiling (**Fig. 2D & S11A-D)**. CE-MS electropherograms revealed that the characteristic 5-PP-InsP_5_ signal present in WT pupae was largely absent in IP6K RNAi pupae, corresponding to an approximately 70-80% reduction in 5-PP-InsP_5_ levels with almost no or minimal effects on other PP-InsP_5_ isomers (**Fig. 5A & S13D**).

**Figure 5.**
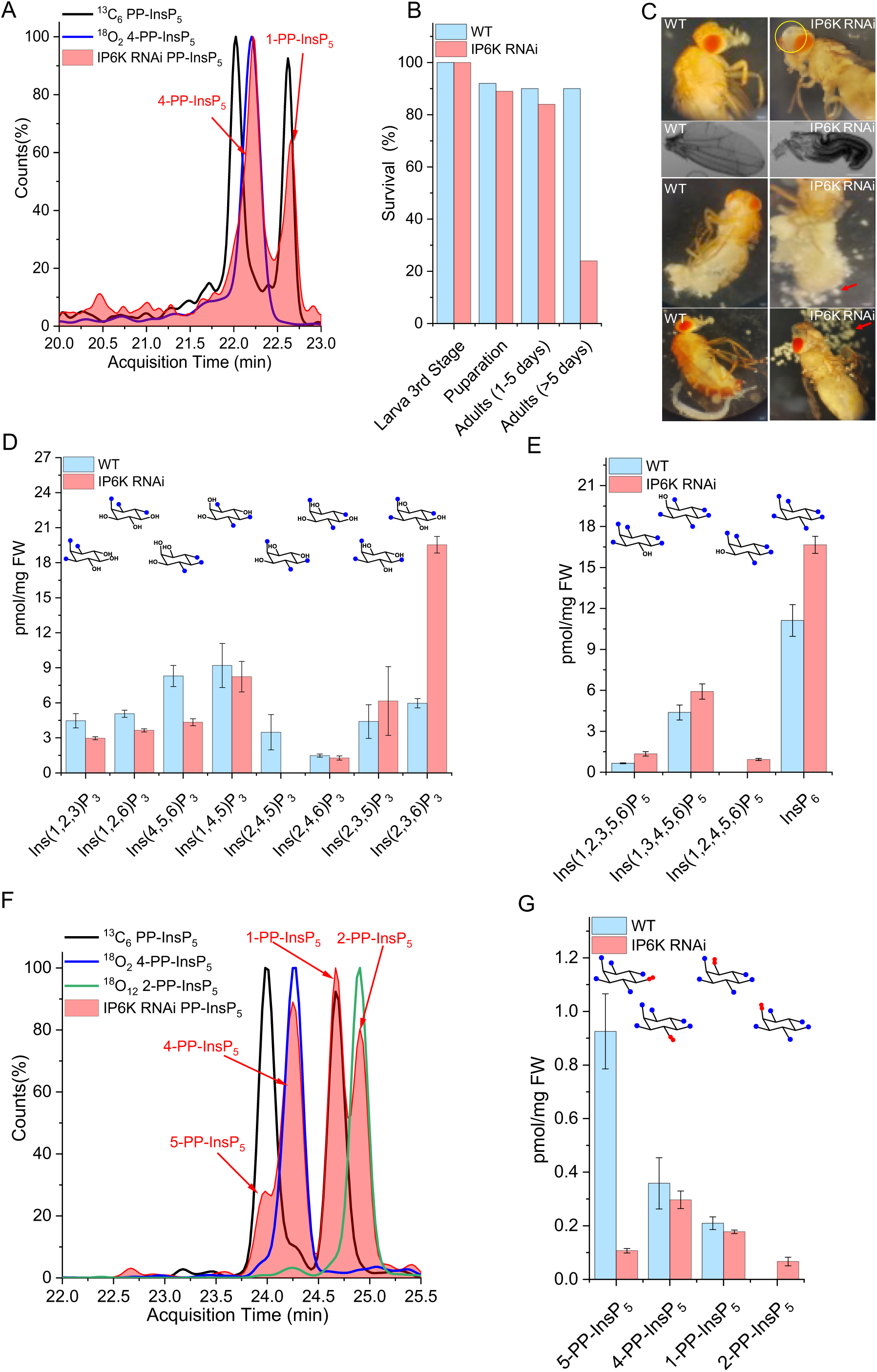
RNAi-mediated depletion of IP6K alters inositol pyrophosphate metabolism and causes post-eclosion physiological defects. (A) Representative CE-MS electropherogram showing separation of PP-InsP_5_ isomers in pupal extracts. The red filled trace corresponds to endogenous PP-InsP_5_ signals, while the black line represents ^13^C-labeled 5-PP-InsP_5_ and 1-PP-InsP_5_ internal references. The blue line represents the ^18^O_2_-labeled 4-PP-InsP_5_ internal reference. Peaks corresponding to 4-PP-InsP_5_ and 1-PP-InsP₅ are assigned with red arrows based on co-migration with the references and accurate mass. (B) Percentage survival of IP6K RNAi flies compared to WT controls. WT shown in grey; IP6K RNAi in red. Although IP6K-depleted flies successfully eclose, 60-70% of adults fail to survive beyond 3 days post eclosion (n=3). (C) Representative adult phenotypes showing normal head, wing, and proboscis morphology in WT flies, whereas IP6K RNAi adults display abnormal head morphology with a prominent dorsal bump-like protrusion, severe wing deformation, impaired proboscis extension, and accumulation of spherical droplet-like structures (indicated with red arrows) within the body cavity. Yellow circle indicate the affected regions. (D) Quantification of InsP_3_ isomers in 12-24 h pupae following RNAi-mediated depletion of IP6K, measured by CE-MS. Bars represent mean ± s.d. (n=3) (WT shown in sky blue; IP6K RNAi in red) and illustrate changes in the InsP_3_ isomer pool. (E) Quantification of InsP_5_ isomers and InsP_6_ in the same samples. Bars represent mean ± s.d. (n=3) (WT in sky blue; IP6K RNAi in red) and show changes in the relative abundance of individual InsP_5_ isomers following IP6K depletion. (F) Representative CE-MS extracted ion electropherogram showing separation of PP-InsP_5_ isomers in pupal extracts. The red filled trace corresponds to endogenous PP-InsP_5_ signals, while the black line represents ^13^C-labeled 5-PP-InsP_5_ (migration time 25.1 min) and 1-PP-InsP_5_ (migration time 25.6 min) internal references. The blue line represents the ^18^O_2_-labeled 4-PP-InsP_5_ and green represents ^18^O_12_-labeled 2-PP-InsP_5_ internal references. (G) Quantification of PP-InsP_5_ isomers in 12-24 h pupae following IP6K depletion. Bars represent mean ± s.d. (n=3) (WT in sky blue; IP6K RNAi in red).

We next assessed developmental phenotypes following RNAi-mediated depletion of IP6K from larva to adult. Although these RNAi IP6K knockdowns proceeded normally through larval development and pupation, a distinct phenotype emerged at the pupal-to-adult transition: >90% adults eclosed but within 3-5 days post-eclosion >70% of these adult flies failed to survive (**Fig. 5B & S17A, Table S3**). Survivors displayed significant morphological anomalies, including unexpanded wings, malformed head structures, and impaired proboscis extension, consistent with widespread failure of tissue maturation (**Fig. 5C & S18A-B**).

5-PP-InsP_5_ was essentially absent in IP6K RNAi 12-24h pupae but they still had levels of 1- and 4-PP-InsP_5_ that were close to wild-type control (**Fig. 5A**). This is in line with IP6K having specific 5-kinase activity. To determine whether loss of 5-PP-InsP_5_ was accompanied by broader changes throughout the InsP network, we examined the distribution of lower InsPs. Unexpectedly, analysis of InsP_3_, revealed a ca. 30% loss of Ins(4,5,6)P_3_ and a complete loss of Ins(2,4,5)P_3_. In contrast, Ins(2,3,5)P_3_ accumulated nearly 3-fold (**Fig. 5D**). At the InsP_4_ level, CE-MS resolved three isomers Ins(1,4,5,6)P_4_, Ins(1,3,4,5)P_4_, and Ins(1,2,5,6)P_4_, together with additional currently not assignable species (**Fig. S13B & S17C**). InsP_5_ isomer composition also shifted markedly, with production of Ins(1,2,4,5,6)P_5_ and enrichment of Ins(1,2,3,5,6)P_5_ accompanied by an overall ca. 40% increase in the total InsP_5_ level (**Fig. 5E**). Also, the absolute InsP_6_ level increased by ca. 30% (**Fig. 5E**). While this increase might be linked to impaired turnover of InsP_6_ to 5-PP-InsP_5_ (**Fig. 5F**), the changes across the whole network indicate broader functions of IP6K.

We also examined adult tissue organization following IP6K depletion. Dissection of IP6K RNAi adults revealed disrupted abdominal fat body organization, with dispersed floating cellular clusters replacing the normally compact adipose tissue (**Fig. 5C & S18A-D**). This phenotype suggests a severe failure of larval fat body cells to remodel successfully into their adult counterparts during pupal development and metamorphosis. In addition, adult head structures did not form properly, suggesting morphogenetic events during head metamorphosis and eversion did not proceed properly. Consequently, emerging adult flies survived for only a few days. We also profiled InsP network alterations in surviving 3-5 day old IP6K RNAi adults. Consistent with pupal observations, InsP_6_ levels were increased by ca. 20%, whereas 5-PP-InsP_5_ and 1,5-(PP)_2_-InsP_4_ were nearly undetectable. Notably, an additional prominent 2-PP-InsP_5_ signal appeared as previously found in the IPK2 knockdown, potentially pointing towards an interplay of the two kinases. Isomer identity was assigned by co-migration with a synthetic ^18^O_12_-labeled 2-PP-InsP_5_ internal reference as described above (**Fig. 5G**).

## Discussion

With the advent of mass spectrometry-based methods in InsP and PP-InsP profiling that do not require radioactive isotope labeling, organs and even whole organisms have become available for detailed analyses^54^. Here, we provide the first isomer-resolved, in-depth analysis of the InsP and PP-InsP network in *Drosophila melanogaster* across its life cycle. Our analysis demonstrates that development is accompanied by changes in a vast InsP network whose reconfiguration parallels morphological and physiological transitions. High-resolution CE-MS analyses with stable isotope ^13^C- and ^18^O-labeled standards, enabled positional isomer assignments across developmental stages (**Fig. S19**). The analysis also reveals that InsP flux is dynamically remodeled rather than constitutively maintained. At specific developmental transitions, the production of defined isomers becomes critical, suggesting that stage-specific InsP network composition contributes to accurate progression. We therefore included three key enzymes in our analyses that regulate this network: IPK1, IPK2, and IP6K.

A first notable feature of *Drosophilás* InsP network is the unusually high diversity of lower inositol phosphates during early development, exceeding even that of InsP_3_ isomers previously identified in duckweed^42^. Egg to larval transition contain multiple non-canonical InsP_3_ isomers, including Ins(1,2,6)P_3_, Ins(2,3,5)P_3_, and Ins(2,4,5)P_3_ (**Fig. 2A**). These do not derive directly from PtdIns(4,5)P₂ hydrolysis^72^ and therefore a complex enzymology in *D. melanogaster* can be anticipated. The range of InsP_3_ and InsP_4_ isomers observed here exceeds by far what has typically been reported in any eukaryotic system. For example, CE-MS analyses in budding yeast, human HCT-116 cells, and human urine have generally resolved fewer InsP_3_ isomers dominated by Ins(1,2,3)P_3_ and Ins(1,2,6)P_3_^46,47^.

Importantly, this diversity lasts during late larval stages and pupation, after which the isomeric pattern of InsP_3_-InsP_4_ progressively converges towards a more restricted profile with accumulation of Ins(1,4,5)P_3_, Ins(4,5,6)P_3_ and its downstream metabolites, e.g. Ins(1,3,4,5)P_4_, Ins(1,3,4,5,6)P_5_, InsP_6_ and PP-InsPs (**Fig. 2A**). This may indicate a progressive activation of phospholipase C signaling and an increasing flow through the lipid-dependent InsP pathway^73^. This change from early isomeric diversity to a canalized adult metabolic profile suggests that early developmental stages may rely on a wider InsP isomer pool and multiple pathways to support growth and tissue patterning, while metamorphosis then needs a more restricted pathway organization that supports stable physiological function.

Genetic perturbation of key enzymes demonstrates the importance of the functioning network. IPK2 depletion prevents the developmental transition towards canonical InsP_4_ and InsP_5_ species (**Fig. 3 & S14)**, and strongly reduces InsP_6_ (**Fig. 3**). These metabolic disruptions are likely involved in the significant developmental arrest previously reported in IPK2 mutants, which do not achieve metamorphosis due to compromised imaginal-disc proliferation and increased apoptosis^6^. Together, these findings suggest that IPK2 serves as a crucial metabolic hub for establishing the higher-order InsP network.

IPK1 depletion remodels the InsP_4_ pool, leading to accumulation of Ins(1,4,5,6)P_4_ and related species (**Fig. S14E**). Because Ins(1,4,5,6)P_4_ has been shown to function in chromatin remodeling and transcriptional regulation in mammals^3,65,74,75^, its accumulation raises the possibility that InsP_4_ signaling may be contributing to transcriptional reprogramming during metamorphosis. In addition, IPK1 depletion results not only in the accumulation of InsP_5_ but selectively enriches atypical isomers like Ins(1,2,4,5,6)P_5_ and Ins(1,2,3,4,6)P_5_, indicating redistribution of flux within the InsP_5_ pool (**Fig 4B-C**). Previous studies showed that loss of IPK1 activity alters inositol phosphate metabolism, although earlier methods lacked the ability to assign positional isomers and measured total InsP_5_ levels^2,7,73^. Together, these observations support the role of *Drosophilás* IPK2 and IPK1 in channeling diverse lower-order intermediates into a restricted pathway leading to InsP_6_ production. However, the enzymatic origin of atypical isomers remains unclear, raising the possibility of additional kinase activities or alternative metabolic routes contributing to InsP_5_ diversification.

PP-InsPs undergo especially dramatic changes during development. The main likely product of *Drosophila* IP6K, 5-PP-InsP_5_, rises sharply in late larval and pupal stages, and becomes the dominant pyrophosphate in adults (**Fig. 2D**). In contrast, 1-PP-InsP_5_ and 4-PP-InsP_5_ are enriched during early developmental stages, suggesting that multiple pyrophosphate species contribute to overall PP-InsP signaling. Similar positional PP-InsP_5_ isomers have also been detected in mammalian systems and plants using CE-MS^47,49,76^, but this study provides the first report of 4-PP-InsP_5_ in fruit flies. RNAi-mediated depletion of IP6K results in near-complete loss of 5-PP-InsP_5_, while other PP-InsP_5_ isomers remain largely unchanged (**Fig. 5G**), supporting the notion that IP6K functions as the physiological 5-kinase. Despite relatively normal metamorphosis, IP6K-depleted *D. melanogaster* exhibit defects in fat-body maturation, abnormal morphology, and significantly reduced adult survival (**Fig. 5C, S18**).

Surprisingly, knockdown of both IP6K and IPK2 led to the production of another PP-InsP_5_ isomer: 2-PP-InsP_5_ (**Fig. 3C, 5F**). While the presence of this metabolite in mammals has been suggested before^45^, it has not been unambiguously proven with a heavy isotope labeled reference as provided in this study. The striking emergence of 2-PP-InsP_5_ as a consequence of kinase knockdown highlights the plasticity of the PP-InsP network. It suggests that disruption of certain kinases affects substrate channeling by protein-protein interactions that alter specificity or that certain kinases also function in the reverse way as phosphatases. Such functional promiscuity has been described for e.g. plant ITPK^77^. The loss of IPK1 also led to disappearance of 4-PP-InsP_5_ and strong reduction of 1-PP-InsP_5_ (**Fig. 4D**). Could IPK1 therefore host additional kinase activity capable of producing inositol pyrophosphates? Again, such flexibility has been described previously in plants, where ITPK enzymes can generate 5-PP-InsP_5_, whereas IPMK-like kinases produce 4-PP-InsP_5_^18^. Overall, these observations raise several interesting mechanistic questions for future studies.

Taken together, our results provide a detailed picture of the *Drosophila melanogaster* inositol phosphate pathway, with IPK2, IPK1, and IP6K acting as key enzymes that regulate InsP flux during development (**Fig. 6**). Early development is marked by an unprecedented array of lower InsP isomers, while metamorphosis reduces this initial complexity. *D. melanogaster’s* most abundant inositol phosphate is InsP_6_, which can be transformed into pyrophosphorylated derivatives (**Fig. 6**). One key enzyme in this pathway is IP6K, but it is apparent that other kinases must also generate additional PP-InsPs. We thus expect that new enzymatic activities can be discovered in the *Drosophila melanogaster* model. Future studies should now address the discovered InsP diversity and connect it to specific functions during development and how the network (and its changes) may coordinate signaling and morphogenesis throughout the *Drosophila melanogaster* life cycle.

**Figure 6.**
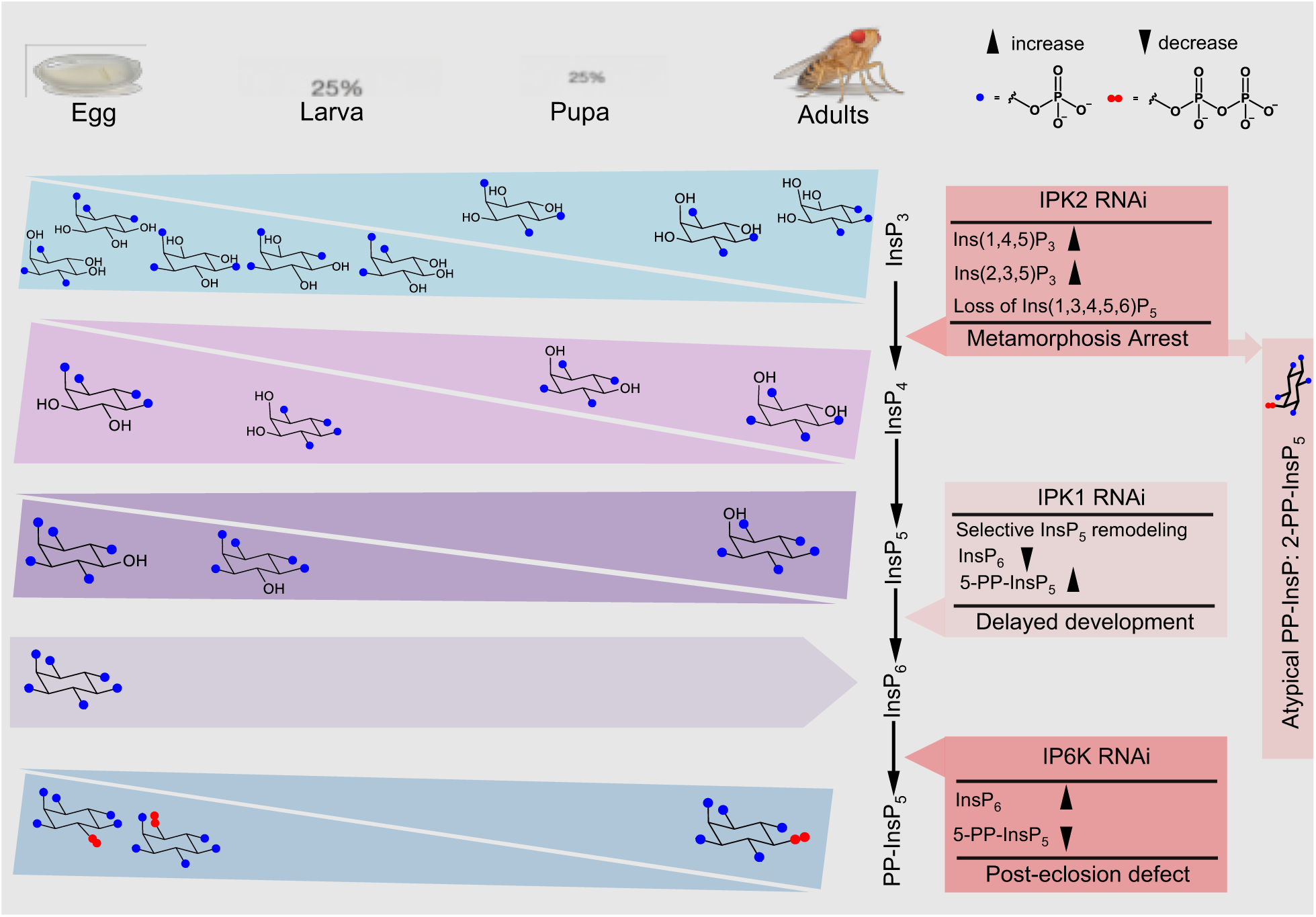
Overview of developmental changes of the inositol phosphate network in *Drosophila melanogaster*. Schematic representation of stage-specific remodeling of the inositol phosphate (InsP) network across egg, larval, pupal, and adult stages. Early development is characterized by a broad diversity of InsP_3_ and InsP_4_ positional isomers present at relatively low abundance, indicating a metabolically flexible network. During larval-to-pupal transition, this diversity progressively resolves into a pathway dominated by canonical intermediates, including Ins(1,3,4,5,6)P_5_ and InsP_6_. In parallel, the inositol pyrophosphate pool shifts from early dominance of 1-PP-InsP_5_ and 4-PP-InsP_5_ towards 5-PP-InsP_5_ in later stages. The red boxes indicate effects of RNAi-mediated depletion of core kinases. IPK2 depletion affects InsP_4_ and InsP_5_ production, leading to loss of canonical InsP_5_ species, and developmental arrest at metamorphosis. IPK1 depletion results in selective InsP_5_ changes, reduced InsP_6_, and delayed development. IP6K depletion results in decrease of 5-PP-InsP_5_, which impairs adult maturation. Notably, despite being kinase knockdowns, they induce the emergence of a non-canonical PP-InsP_5_ species, identified as 2-PP-InsP_5_.

## Methods

### *Drosophila melanogaster* maintenance, developmental staging and RNAi

All experiments employed *Drosophila melanogaster*, with fly strains specified in Supplementary **Table S1**. Flies were kept in a standard cornmeal-yeast-agar medium at 18–22 °C under a controlled laboratory setting. For developmental profiling, synchronized wild-type flies were generated by timed egg laying. Samples were collected at defined development stages including eggs, larvae, pupae, and adults, flash-frozen and stored at −80 °C until extraction.

RNAi-mediated gene knockdown was performed using UAS-driven hairpin constructs obtained from the Vienna Drosophila Resource Center (VDRC). RNAi induction was achieved using a heat-shock–inducible FLP/GAL4 system^78^. Briefly, crosses were established using the hsFlp[122]; tub-GAL4, UAS-mCD8::GFP driver line to enable inducible GAL4 activation following heat shock. This strategy allows temporal control of RNAi expression during development while maintaining a consistent genetic background. GFP expression served as a marker for GAL4 activation. Further details can be found in the supplementary methods.

### Inositol phosphates extraction, TiO_2_ enrichment, and CE-MS analysis

Flash-frozen whole *Drosophila* samples from different stages, were homogenized in chilled 1 M perchloric acid followed by gentle rotation at 4 °C and centrifuged to obtain soluble inositol phosphates metabolites. To enrich inositol phosphates and pyrophosphates from these extracts, we used TiO_2_ bead-based affinity purification^55^. The dried samples were then stored at -20 °C, until reconstitution and analysis by CE-MS.

CE-MS analyses were performed with an Agilent 7100 CE system coupled to either a Triple Quadrupole (QqQ) or quadrupole time-of-flight (QToF) mass spectrometer operating in negative ionization mode. Inositol phosphate species were separated using ammonium acetate-based background electrolytes optimized for either general profiling or enhanced positional isomer resolution^45,49,50^. Confident positional assignment and quantification of inositol phosphate isomers were achieved using a complementary dual-isotope strategy involving ^13^C_6_- and ^18^O_12_-labeled standards. Further details are provided in the supplementary methods.

## Acknowledgements

A.S. acknowledges CIBSS for a Post-Doctoral Fellowship. A.S. thanks Ananthakrishnan Vijayakumar Maya, Lara Heckmann and the staff of the Life Imaging Centre (LIC) in the Hilde Mangold House (HMH) of the Albert-Ludwigs-University of Freiburg for help with their microscopy resources. A.S. thanks Isabelle Grass for helping during fly collections. We thank Gabriel Schaaf for critically proofreading the manuscript. This study was supported by the Deutsche Forschungsgemeinschaft (DFG, Project JE 572/11-1) and under Germany’s Excellence Strategy (CIBSS, EXC-2189, Project ID 390939984, to H.J.J; A.K.C). H.J.J. acknowledges funding from the Volkswagen Foundation (VW Momentum Grant 98604).

## Contributions

A.S., A.K.C., and H.J.J. designed the experiments and wrote the manuscript. A.S. performed the experiments and wrote the original draft. M.B. and M.H. synthesized the compounds. A.S., M.B., A.K.C. and H.J.J. reviewed and edited the manuscript draft.

