## Supplemental Information for "The dynamic inositol phosphate network during development in *Drosophila melanogaster*"

**Supplementary Information**

**Supplementary Results and Discussion:	p. 2**

**Supplementary Tables:	p. 5**

**Supplementary Figures:	p. 8**

**Supplementary Methods:	p. 28**

**Supplementary References:	p. 32**

**Supplementary Results and Discussion**

**Preparation of *Drosophila* Samples for CE–MS Analysis**

To enable accurate comparison of inositol phosphate levels across developmental stages, all measurements were normalized to the fresh (wet) weight of the starting material. This normalization strategy was selected because *Drosophila melanogaster* undergoes substantial stage-dependent changes in tissue composition, water content, and overall physiology during development, particularly throughout larval growth and metamorphosis. Under these conditions, protein content does not provide a reliable basis for normalization of whole-organism metabolite data. Previous studies have similarly reported pronounced developmental fluctuations in protein abundance that primarily reflect tissue remodeling and organismal growth rather than proportional changes in metabolite levels^1,2^. Protein concentration does not consistently indicate the organism's size or metabolic activity at each developmental stage, rendering it unsuitable for normalizing metabolomics data during development. We normalized to fresh tissue mass as recommended in metabolomics studies^3–6^. This approach is commonly applied to whole organisms or tissues, especially in systems where cell number, protein content, and water composition change substantially during development and therefore cannot be reliably normalized between samples. Samples were weighed and immediately processed for metabolite extraction. For each biological replicate, individual flies were pooled to reduce biological variability and to obtain sufficient material for analysis. Therefore, all quantitative CE–MS data in this study are reported relative to fresh tissue mass, enabling robust and reliable comparisons of inositol phosphate levels across developmental stages.

**Separation and stereochemical assignment of inositol isomers by CE–MS**

Free inositol and all phosphorylated inositol species identified in this study are presumed to originate from the *myo*-inositol configuration, which serves as the canonical and biologically significant scaffold in eukaryotic systems^7^. However, since inositol can occur in principle in several stereoisomeric forms we initially assessed, which inositol species predominates in *Drosophila*. Apart from the *myo*-configuration, the *scyllo*-configuration has been described as a common inositol species, particularly in soils. We therefore studied these two species as potential constituents of the inositol network in *Drosophila*. Baseline separation of *myo*- and *scyllo*-inositol standards was first established using CE-MS under the same analytical conditions (**Table S5**) applied for inositol phosphate profiling (**Fig. S2 A–C**). The migration times of the standards were determined individually (**Fig. S2A, B**). For matrix comparability with biological samples, standards were prepared in 1 M perchloric acid, the same extraction medium used for *Drosophila* samples. Co-injection experiments showed baseline separation of the two stereoisomers (**Fig. S2C**). For biological analysis, approximately 300 mg of adult *Drosophila melanogaster* tissue was extracted in ice-cold 1 M perchloric acid followed by TiO_2_ enrichment as described in the main Methods. Free inositol was not found in the enriched extracts, as TiO_2_ does not retain unphosphorylated metabolites. Samples were then digested either with BASF Natuphos or heated in acidic condition (2M per chloric acid, pH ≥ 0) at 95 °C for 12 h (**Fig. S2D, E**) for enzymatic and thermal dephosphorylation of inositol phosphates and release of free inositol prior to CE–MS analysis. Such treatment resulted in one major and one minor signal. Based on the standard separation, the minor peak was assigned to *scyllo*-inositol (even though we note that it could be other forms of inositol that potentially comigrate). To confirm this assignment, heat-digested extracts were spiked with 2 µM *scyllo*-inositol and re-analyzed under the same CE–MS conditions (**Fig. S2F**). The detected peak comigrated with *scyllo*-inositol. All electropherograms are extracted ion signals recorded under the same CE–ESI–MS conditions.

**TiO_2_ based InsPs recovery**

To evaluate the efficiency of inositol phosphate extraction and TiO_2_-based enrichment, recovery control experiments were performed using biological samples processed identically to those used in the main study. Known amounts of isotopically labeled standards - 2.5 µM [^13^C_6_]-InsP_6_, and [^13^C_6_]-Ins(1,3,4,5,6)P_5_ - were added either prior to extraction (pre-spiking) or after extraction immediately before CE–MS analysis (post-spiking). Samples were processed as described in the extraction methodology. Recovery efficiencies were determined by comparing the CE–MS signals (peak areas) obtained from pre-spiked ^13^C_6_-labeled standards in the samples with those from post-spiked ^13^C_6_-labeled standards in the samples, thereby accounting for analyte losses occurring during extraction and enrichment. Using this approach, average recovery rates of 79% for InsP_6_, and 72% for Ins(1,3,4,5,6)P_5_ were obtained (**Fig. S3**). Values represent the mean of three independent experiments.

**Synthesis of ^18^O_6_-labeled Ins(2,4,6)P_3_**

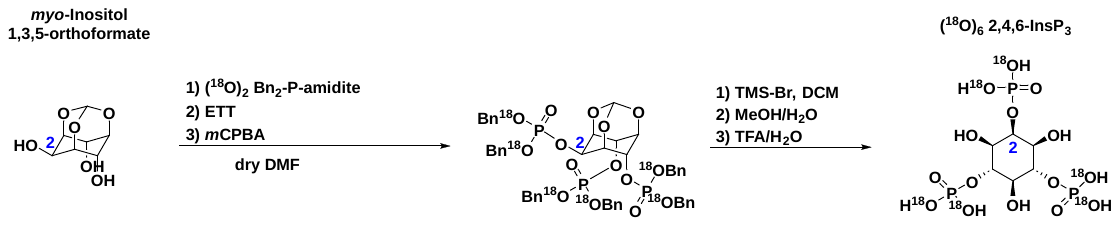

**General Remarks**

Semi-preparative RP-HPLC was performed using a PuriFlash 5.250P by Advion Interchim Scientific. The stationary phase consists of a PF5C18AQ-150/100 Prep-LC column supplied by Advion Interchim Scientific.

Lyophilizations were done with a Christ Freeze Dryer Alpha 1-4 LDplus.

NMR spectra were recorded on a Bruker Avance 400 MHz spectrometer in the indicated deuterated solvent. Data are reported as follows: chemical shift (δ, ppm), multiplicity (s: singlet, d: doublet, t: triplet; q: quartet; m: multiplet; ψ: pseudo-multiplet), coupling constant(s) (J, Hz), integration, assignment. All signals were referenced to the internal solvent signal as standard (D_2_O: δ = 4.79 ppm, CDCl_3_: δ = 7.26 ppm). Deuterated solvents were obtained from commerical suppliers (Euroisotop, Deutero) and used without further purification.

Mass spectra were recorded by the mass spectrometry service of the University of Freiburg.

**(1*R*,3*s*,5*r*,6*R*,7*S*,8*s*,9*S*)-2,4,10-Trioxaadamantane-6,8,9-triyl hexabenzyl tris(phosphate-^18^O_2_)**

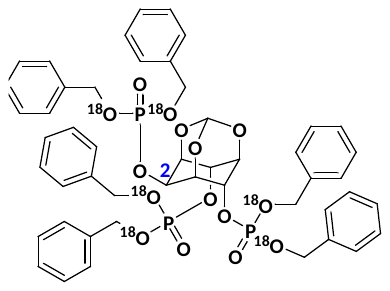

A flame-dried Schlenk tube was charged with *myo*-inositol 1,3,5-orthoformate (12.0 mg, 63.1 μmol, 1.0 eq.) and (^18^O)_2_ Bn_2_-P-amidite (132 mg, 129 μL, 379 μmol, 6.0 eq.) and co-evaporated with dry acetonitrile (1 ml) twice. The mixture was dissolved in dry DMF (1.5 ml) and ETT (49.3 mg, 379 μmol, 6.0 eq.) was added. The reaction mixture was stirred at room temperature for 1 h, cooled down to 0°C and *m*CPBA (70%, 93.3 mg, 379 μmol, 6.0 eq.) was added. After stirring for 10 min the reaction was allowed to warm to room temperature and was stirred for further 10 min.

For purification, the mixture was directly subjected to automated reversed-phase semi-preparative HPLC (Interchim PF5C18AQ-150/100 column, H_2_O/MeCN, gradient: 0-90% MeCN, 10% triethylammonium acetate buffer (0.1 M)). The product *myo*-inositol 2,4,6-(OP(O)^18^O_2_Bn_2_)_3_ 1,3,5-orthoformate was obtained as a colorless solid (46.0 mg, 46.8 μmol, 74%).

**HRMS (ESI+)** m/z for C_49_H_50_O_9_^18^O_6_P_3_ [M+H]^+^: calcd. 983.2612, found 983.2633.

NMR spectra were in accordance with the non ^18^O-labeled analog from J.-M. Lehn, S. Pothukanuri, A. E. Koumbis, C. Duarte, N. Claude, “Cyclitols and their Derivatives and their Therapeutic Applications”, 2008, WO 2008/082658 Α2.

**[^18^O_6_] Ins(2,4,6)P_3_**

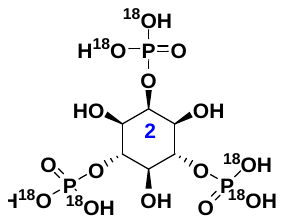

In a flame-dried Schlenk tube *myo*-inositol 2,4,6-(OP(O)^18^O_2_Bn_2_)_3_ 1,3,5-orthoformate (46.0 mg, 46.8 μmol, 1.0 eq.) was dissolved in dry DCM (3 mL). Then TMS-Br (129 mg, 111 μL, 842 μmol, 18.0 eq.) was added and the reaction mixture was stirred at room temperature under argon overnight. The reaction mixture was concentrated under reduced pressure at 40°C and re-dissolved in methanol (3 mL) and water (1 mL). The reaction mixture was stirred at room temperature for 1 h and the intermediate precipitated by addition of diethyl ether (‑20°C). The precipitate was re-dissolved in 2% TFA in H_2_O (2 mL) and stirred at room temperature for 3 days. The reaction solution was evaporated under reduced pressure at 40°C.

The solid was re-dissolved in water and subjected to ion-exchange resin column chromatography (Chelex 100 sodium form). The product [^18^O_6_] 2,4,6-InsP_3_ was obtained as a colorless solid (13.2 mg, 23.4 μmol, 50%) in form of the sodium salt.

**^31^P{^1^H}-NMR** (162 MHz, D_2_O, pH=1): δ 0.12 (s, 2P, P-4, P-6), 0.04 (s, 1P, P-2) ppm.

**^1^H-NMR** (400 MHz, D_2_O, pH=1): δ 4.62 (dψt, J = 8.6, 2.6 Hz, 1H, H-2), 4.24 (ψq, J = 9.3 Hz, 2H, H-4, H‑6), 3.76 (dt, J = 9.9, 2.2 Hz, 2H, H-1, H-3), 3.60 (ψt, J = 9.2 Hz, 1H, H-5) ppm.

**^13^C{^1^H}-NMR** (101 MHz, D_2_O, pH=1): δ 78.7 (d, J = 6.2 Hz, C-4, C-6), 78.4 (d, J = 6.3 Hz, C-2), 72.3 (t, J = 3.5 Hz, C-5), 69.4 (t, J = 3.3 Hz, C-1, C-3) ppm.

**HRMS (ESI-)** m/z for C_6_H_14_O_9_^18^O_6_P_3_ [M-H]^-^: calcd. 430.9806, found 430.9813.

See **Fig. S5F** for ^1^H-NMR of (^18^O)_6_ 2,4,6-InsP_3_ (D_2_O, 400 MHz, pH = 1).

See **Fig. S5G** for ^13^C{1H}-NMR of (^18^O)_6_ 2,4,6-InsP_3_ (D_2_O, 101 MHz, pH = 1).

See **Fig. S5H** for ^31^P{^1^H}-NMR of (^18^O)_6_ 2,4,6-InsP_3_ (D_2_O, 162 MHz, pH = 1).

**Supplementary Tables**

**Table S1. *Drosophila* fly lines used in this study.** Listed are the control, RNAi, and driver lines used for genetic perturbation of inositol phosphate pathway enzymes. Potential RNAi off-target or insertion-site considerations are indicated where relevant.

| **S.No** | **Category** | **Target gene** | **Fly line/genotype** | **Stock ID** | **Source** | **Chromosome** |
| --- | --- | --- | --- | --- | --- | --- |
| **1.** | Control |  | w[1118] | - | Lab stock | - |
| **2.** | Driver |  | hsflp [122]; Sp/CyO, ubi-GFP; UAS-mCD8::GFP, tub-GAL4/TM6c |  |  | - |
| **3.** | RNAi | IPK1 | UAS-IPK1 RNAi | KK-109497 | VDRC KK | Chr 2 |
| **4.** |  | IPK2 (IPMK) | UAS-IPK2 RNAi | GD-43825 | VDRC GD | Chr 2 |
| **5.** |  | IP6K | UAS-IP6K RNAi | KK-103749 | VDRC KK | Chr 2 |
| **6.** |  | IP6K | UAS-IP6K RNAi | GD-38327 | VDRC GD | Chr 2 |

* KK and GD lines were obtained from the Vienna Drosophila Resource Center (VDRC)

**Table S2. Replicate for the Stage specific InsPs isomer**

See Excel file Table S2

**Table S3. Developmental progression and adult survival following RNAi-mediated depletion of IP6K.**

Third-instar larvae (n = 100 per batch) were scored for pupariation, adult eclosion (1–5 days), and survival beyond 5 days after eclosion.

|  |  | **Larva 3rd Stage** | **Pupation** | **Adults eclosed (1-3 days)** | **Adults eclosed (more than 5 days)** |
| --- | --- | --- | --- | --- | --- |
| **WT** | **Batch 1** | **100** | **95** | **93** | **93** |
|  | **Batch 2** | **100** | **90** | **90** | **90** |
|  | **Batch 3** | **100** | **92** | **89** | **89** |
| **IP6K RNAi** | **Batch 1** | **100** | **89** | **85** | **27** |
|  | **Batch 2** | **100** | **85** | **83** | **19** |
|  | **Batch 3** | **100** | **93** | **88** | **23** |
|  | **Batch 4** | **100** | **90** | **84** | **30** |
|  | **Batch 5** | **100** | **88** | **83** | **22** |

**Table S4. Instrument and scan source parameters of QqQ Mass Spectrometer**

| **Parameter** | **Setting** |
| --- | --- |
| Gas temperature | 150 °C |
| Gas flow | 11 L min^-1^ |
| Nebulizer pressure | 8 psi |
| Sheath gas temperature | 175 °C |
| Sheath gas flow | 8 L min^-1^ |
| Capillary voltage | −2000 V |
| Nozzle voltage | 2000 V |
| High-pressure RF (ion funnel) | 70 V |
| Low-pressure RF (ion funnel) | 40 V |

**Table S5. MRM transition settings of InsPs for *Drosophila* samples**

| **Molecular name** | **Precursor ion (m/z)** | **Product ion (m/z)** | **Dwell (ms)** | **Fragmentor (V)** |  | **Collision energy (V)** | **Cell accelerator (V)** | **Polarity** |
| --- | --- | --- | --- | --- | --- | --- | --- | --- |
| ^13^C_6_-ATP | 516 | 417.9 | 110 | 166 |  | 25 | 1 | Negative |
| ATP | 506 | 407.9 | 200 | 166 |  | 25 | 1 | Negative |
| Ins | 179 | 161 | 80 | 166 |  | 25 | 1 | Negative |
| InsP_3_ | 418.9 | 320.8 | 50 | 166 |  | 17 | 4 | Negative |
| ^18^O_6_-InsP_3_ | 424.9 | 326.8 | 50 | 166 |  | 17 | 4 | Negative |
| InsP_4_ | 499.0 | 418.9 | 50 | 166 |  | 5 | 1 | Negative |
| ^18^O_8_-InsP_4_ | 505.0 | 424.9 | 50 | 166 |  | 5 | 1 | Negative |
| InsP_5_ | 579.0 | 498.9 | 50 | 166 |  | 9 | 3 | Negative |
| ^13^C_6_-InsP_5_ | 585.0 | 504.9 | 50 | 166 |  | 9 | 3 | Negative |
| ^18^O_10_-InsP_5_ | 585.0 | 504.9 | 50 | 166 |  | 9 | 3 | Negative |
| InsP_6_ | 659.0 | 480.9 | 50 | 166 |  | 13 | 4 | Negative |
| ^13^C_6_-InsP_6_ | 665.0 | 486.9 | 50 | 166 |  | 13 | 4 | Negative |
| PP-InsP_5_ | 739.0 | 319.9 | 50 | 166 |  | 9 | 3 | Negative |
| ^13^C_6_-5-PP-InsP_5_ | 745.0 | 322.9 | 50 | 166 |  | 9 | 3 | Negative |
| ^18^O_2_-2-PP-InsP_5_ | 743.0 | 323.9 | 50 | 166 |  | 9 | 3 | Negative |
| ^18^O_6_-4-PP-InsP_5_ | 751.0 | 327.9 | 50 | 166 |  | 9 | 3 | Negative |
| (PP)_2_-InsP_4_ | 819.0 | 359.8 | 50 | 166 |  | 9 | 1 | Negative |
| ^13^C_6_-(PP)_2_-InsP_4_ | 825.0 | 362.8 | 50 | 166 |  | 9 | 1 | Negative |

**Table S6.** **Isotopic internal standards and final working concentrations for quantitative CE–MS analysis**

| **Isotopic standard** | **Inositol phosphate species** | **Final concentration in sample (µM)** |
| --- | --- | --- |
| ^13^C_6_-ATP | Adenosine triphosphate | 50 |
| ^13^C_6_-Ins(1,2,3)P_3_ | Inositol 1,2,3-trisphosphate | 10 |
| ^13^C_6_-Ins(1,3,4,5,6)P₅ | Inositol pentakisphosphate (2-OH-InsP_5_) | 4 |
| ^13^C_6_-InsP₆ | Inositol hexakisphosphate (InsP_6_) | 8 |
| ^13^C_6_-5-PP-InsP_5_ | 5-diphospho-inositol pentakisphosphate | 1 |
| ^13^C_6_-1-PP-InsP_5_ | 1-diphospho-inositol pentakisphosphate | 0.5 |
| ^13^C_6_-1,5-InsP₈ | 1,5-bis-diphospho-inositol tetrakisphosphate | 0.25 |
| ^18^O_12_-2-PP-InsP_5_ | 2-diphospho-inositol pentakisphosphate | 0.5 |
| ^18^O_6_-4-PP-InsP_5_ | 4-diphospho-inositol pentakisphosphate | 1 |

**Supplementary Figures**

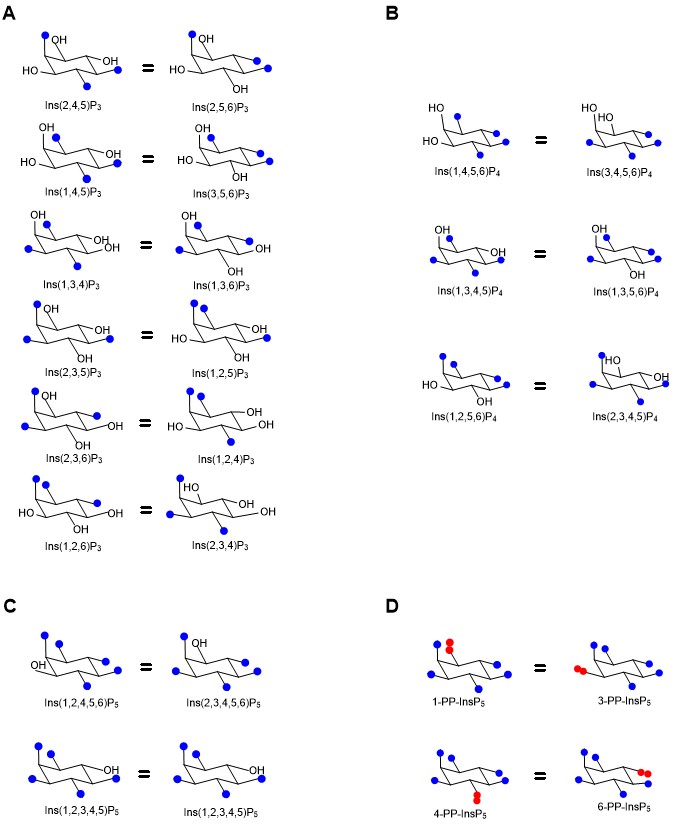

**Supplementary Figure 1. Isomeric and enantiomeric distribution of InsP_3_, InsP_4_, InsP_5_ and PP-InsP_5_ species in *Drosophila melanogaster*.**

Schematic representation of positional isomers detected by CE–ESI–MS for (**A**) InsP_3_, (**B**) InsP_4_, (**C**) InsP_5_ and (**D**) PP-InsP_5_ classes. Blue dots represent phosphate groups and red dots pyrophosphate moieties. Structures correspond to isomers detected in *Drosophila* samples. Enantiomers co-migrate under achiral CE–MS conditions and cannot be distinguished; assignments therefore reflect possible enantiomeric pairs and are reported using lower-numbering convention or the standard reference present.

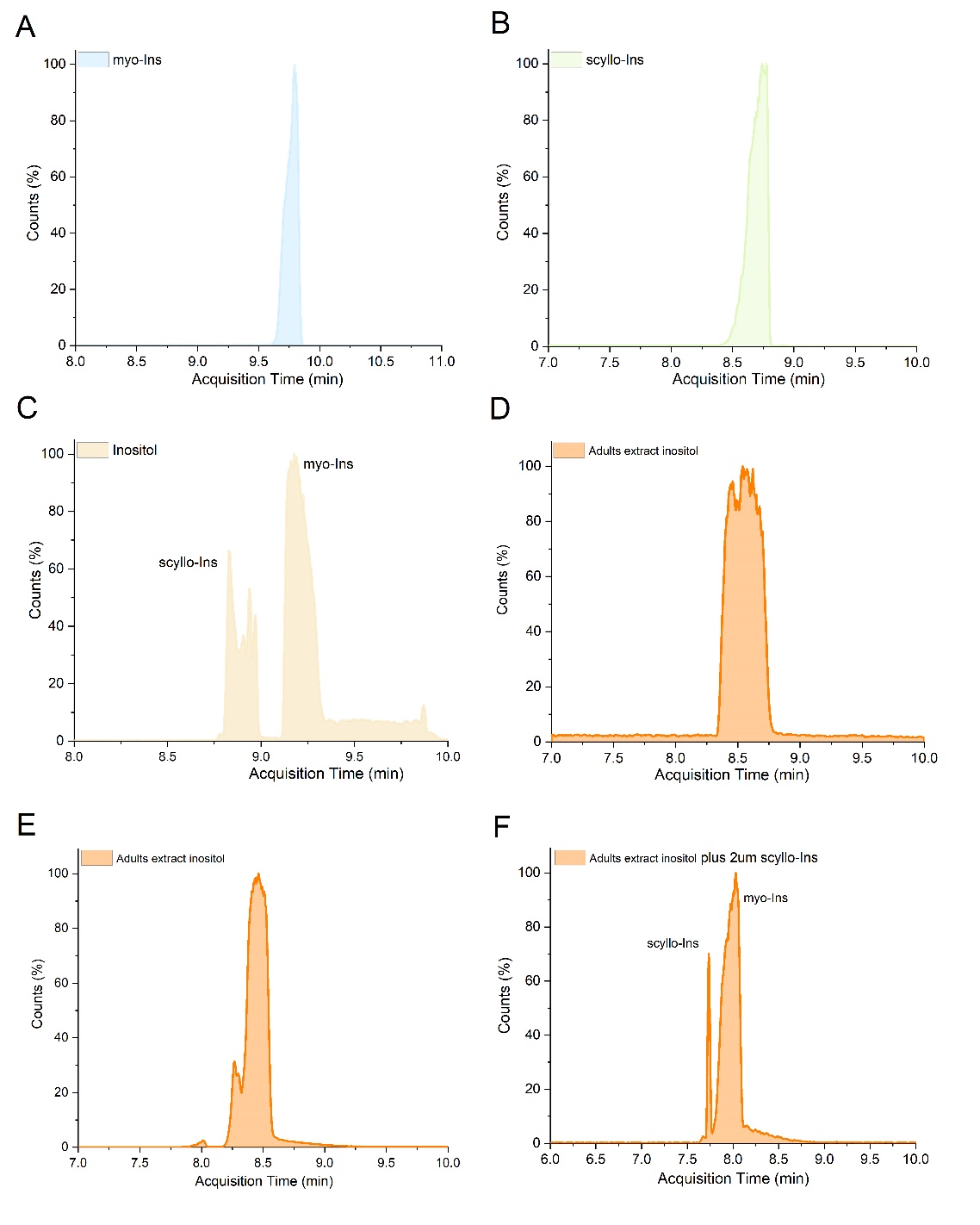

**Supplementary Figure S2. CE–MS separation of *myo*- and *scyllo*-inositol and confirmation of stereochemical identity in adult *Drosophila* extracts.**

**(A)** Extracted ion electropherogram of an authentic **myo-inositol** standard, showing a single signal at its characteristic migration time under the applied CE–MS conditions.

**(B)** Extracted ion electropherogram of an authentic ***scyllo*-inositol** standard, demonstrating a distinct migration time compared to *myo*-inositol.

**(C)** Co-injection of *myo*- and *scyllo*-inositol standards, confirming potential baseline separation between the two stereoisomers under identical analytical conditions.

**(D)** Extracted ion electropherogram obtained from adult Drosophila melanogaster following phytase digestion and release of free inositol.

**(E)** Extracted ion electropherogram obtained from adult *Drosophila melanogaster* extracts following heat-mediated dephosphorylation and release of free inositol.

**(F)** Heat-digested extracts shown in panel E were spiked with 2 µM *scyllo*-inositol and re -analyzed under identical CE–MS conditions. This experiment demonstrates that in this matrix, separation can still be achieved.

Recovery Rate (%)

| Development stages | InsP₆ | Ins(1,3,4,5,6)P₅ |
| --- | --- | --- |
| Egg | 75.9828 | 70.3743 |
| Larva | 82.7586 | 77.308 |
| Pupa | 80.2069 | 71.508 |
| Adults | 80.6897 | 73.21 |

**Supplementary Figure S3. Recovery efficiency of InsP_6_ and Ins(1,3,4,5,6)P_5_ across *Drosophila* developmental stages following TiO₂-based extraction.**

Isotopically labeled standards were used to determine extraction recovery across developmental stages. 2.5 µM of [^13^C_6_]-InsP_6_ and [^13^C_6_]-Ins(1,3,4,5,6)P_5_ were added to biological samples either prior to extraction (pre-spiking) or after extraction but before CE–MS analysis (post-spiking). Recovery efficiency was calculated as the ratio of pre-spiked to post-spiked signal intensities. Recovery rates were consistently similar and high across all stages analyzed, with InsP_6_ recovery ranging from ~76–83% and Ins(1,3,4,5,6)P_5_ recovery ranging from ~70–77% in egg, larval, pupal, and adult samples.

ATP (pmol/mg FW)

| Development stages | pmol/mg |
| --- | --- |
| Egg | 11539.4 |
| Larva | 28241.9 |
| Pupa | 10375.5 |
| Adults | 9609.63 |

**Supplementary Figure S4. Developmental ATP levels in Drosophila melanogaster.**
ATP concentrations were quantified by CE–MS and normalized to fresh (wet) tissue mass across egg, larval, pupal, and adult stages. ATP levels peaked during the larval stage and declined during pupation and adulthood. Data represent mean ± SD from three independent biological replicates.

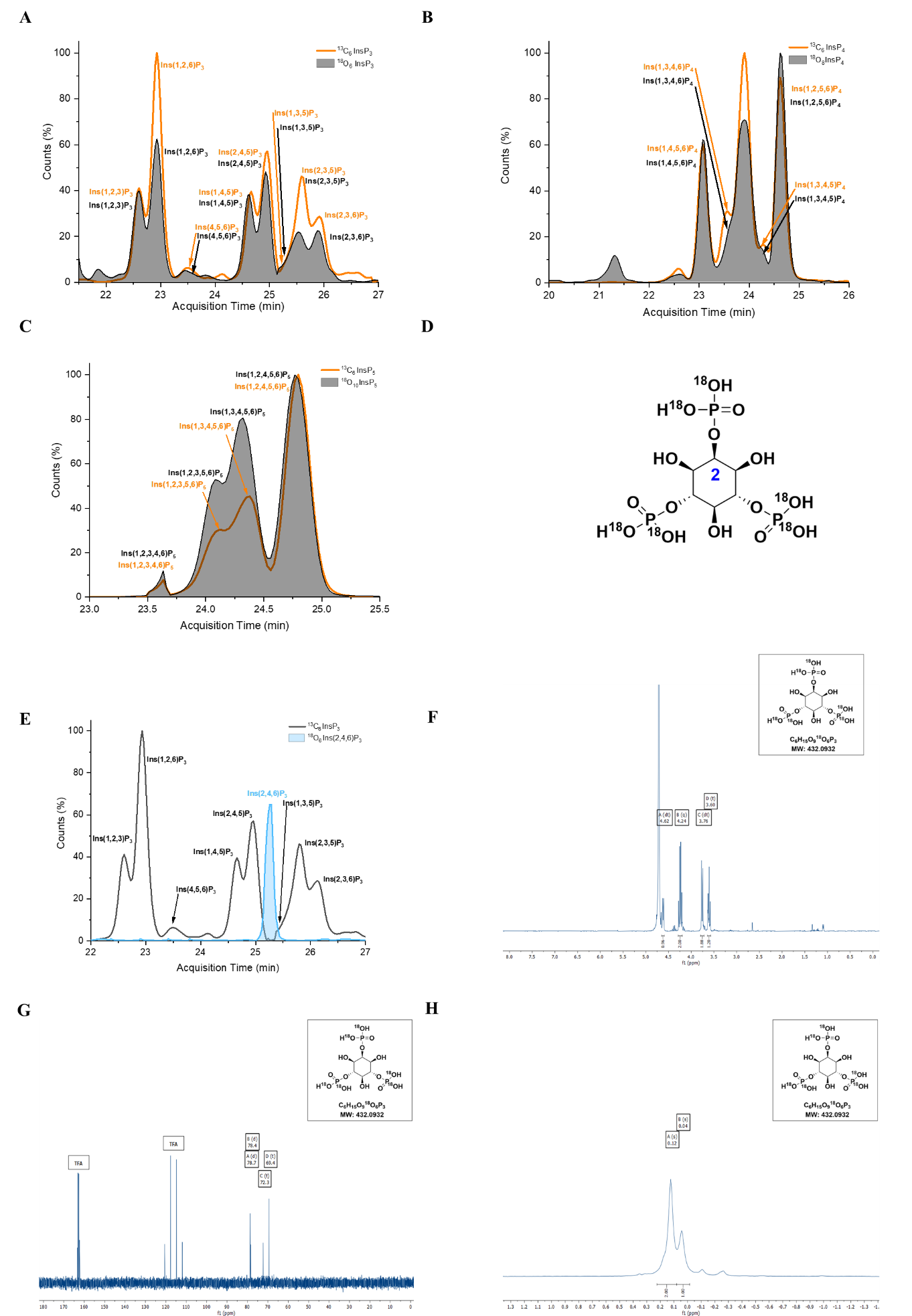

**Supplementary Figure S5. Generation and characterization of isotopically labeled InsP_3_ references for isomer assignment in *Drosophila melanogaster*.**

Extracted ion electropherograms showing comparison of **(A)** [^13^C_6_]-InsP₃ (orange) and [¹⁸O_6_]-InsP₃ (black), (**B**) [^13^C_6_]-InsP_4_ (orange) and [¹⁸O_8_]-InsP₄ (black), (**C**) [^13^C_6_]-InsP_5_ (orange) and [¹⁸O_10_]‑InsP₅ (black) reference mixtures. The [^13^C_6_]-labeled and [^18^O]-labeled InsPs were generated by thermal hydrolysis (pyrohydrolysis) of [^13^C_6_]-InsP_6_ and [^18^O_12_]-InsP_6_ in ultrapure water (100 °C, 5 h). Co-migration of the corresponding [^13^C_6_]- and [^18^O]-labeled InsP_3_, InsP_4_, and InsP_5_ species under identical CE–ESI–MS conditions demonstrates that isotopic substitution does not alter electrophoretic mobility. These [^18^O]-labeled reference mixtures were subsequently used for isomer assignment in *Drosophila* developmental samples. (**D**) Structure of ^18^O_6_-labeled Ins(2,4,6)P_3_ used as a reference standard for metabolite identification. (**E**) CE–ESI–MS electropherograms of InsP_3_ obtained by pyrohydrolysis of ^13^C_6_ InsP_6_ compared to the new ^18^O_6_-Ins(2,4,6)P_3_ show clear separation from the InsP_3_ species, enabling confident isomer assignment with the newly synthesized reference. (**F**) ^1^H-NMR of (^18^O)_6_-2,4,6-InsP_3_ (D_2_O, 400 MHz, pH = 1) (**G**) ^13^C{^1^H}-NMR of (^18^O)_6_-2,4,6-InsP_3_ (D_2_O, 101 MHz, pH = 1) (**H**) ^31^P{^1^H}-NMR of (^18^O)_6_-2,4,6-InsP_3_ (D_2_O, 162 MHz, pH = 1).

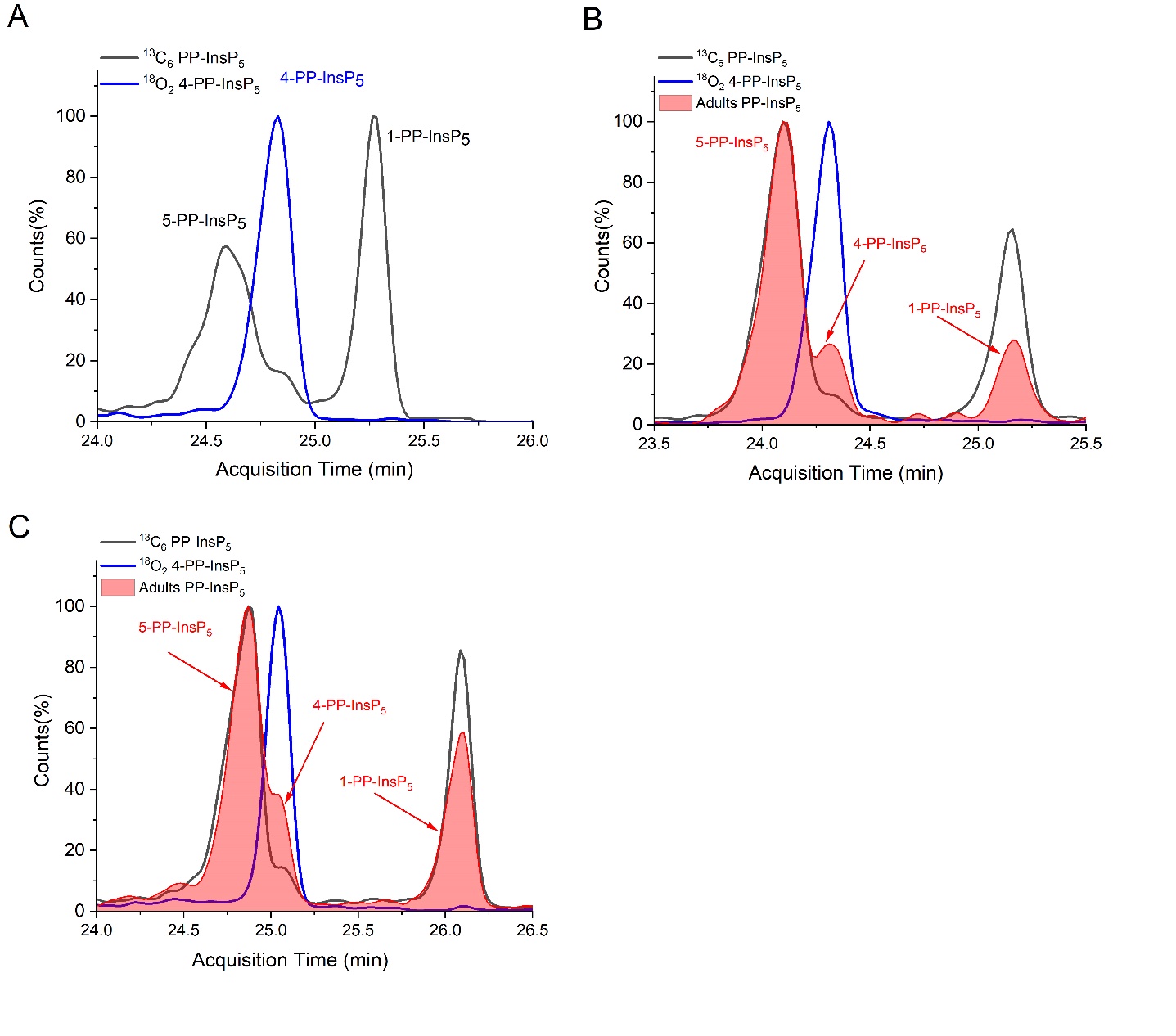

**Supplementary Figure S6. Separation and assignment of PP-InsP_5_ isomers by CE–MS.**

(**A**) CE–MS electropherograms showing separation of PP-InsP_5_ positional isomers, including 5-PP-InsP_5_, 4-PP-InsP_5_ and 1-PP-InsP_5_. (**B,C**) Extraction of 150 mg (**B**) and 300mg (**C**) adult flies with spiked ^13^C_6_ labeled 5-PP-InsP_5_, 1-PP-InsP_5_ and ^18^O_2_ 4-PP-InsP_5_ prior to perchloric acid extraction and TiO_2_ enrichment. Addition of the labeled standards resulted in the expected increase in signal intensity of the corresponding species but did not result in additional PP-InsP_5_ isomers, confirming that little to no isomer interconversion occurs during sample preparation or CE–MS analysis.

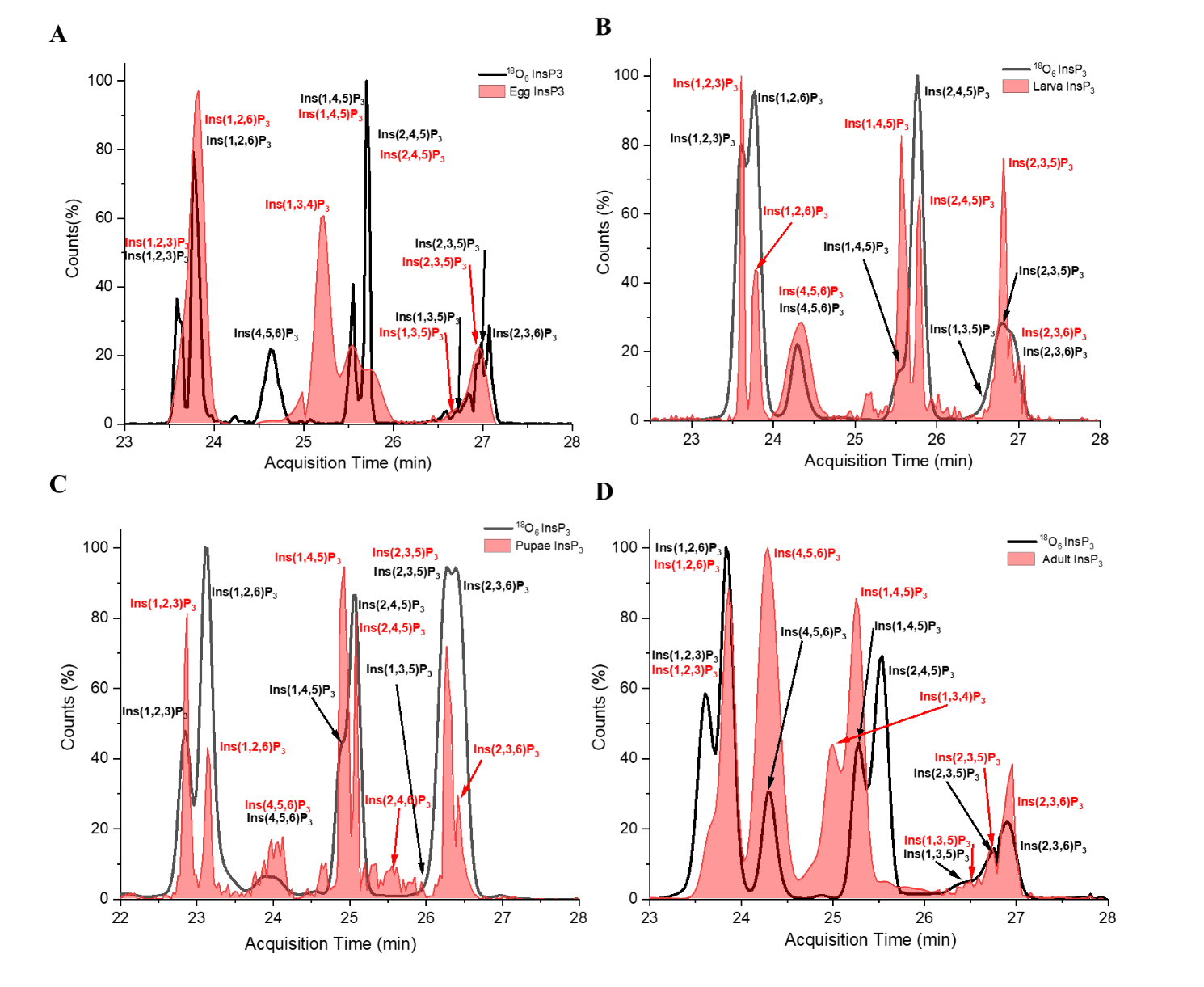

**Supplementary Figure S7. Assignment of InsP_3_ isomers in stage-specific *Drosophila melanogaster* samples using isotopically labelled references.**

Extracted ion electropherograms showing positional assignment of InsP_3_ isomers in stage-specific *Drosophila* melanogaster samples. (**A**) egg, (**B**) larva, (**C**) pupae, and (**D**) adult extracts (red area) are overlaid with an [^18^O_6_]-InsP_3_ reference mixture (black line) analyzed under identical CE–ESI–MS conditions. InsP_3_ isomer assignment was performed by direct comparison of endogenous migration times with those of [¹⁸O₆]-InsP_3_ reference peaks, supported by prior validation using [¹³C₆]-InsP_3_ standards. Endogenous InsP_3_ species co-migrating with [^18^O_6_]-Ins(1,2,3)P_3_, [^18^O_6_]-Ins(1,2,6)P_3_, [^18^O_6_]-Ins(4,5,6)P_3_, [^18^O_6_]-Ins(1,4,5)P_3_, [^18^O_6_]-Ins(2,4,5)P_3_, [^18^O_6_]-Ins(2,4,6)P_3_, [^18^O_6_]-Ins(1,3,5)P_3_, [^18^O_6_]-Ins(2,3,5)P_3_, and [^18^O_6_]-Ins(2,3,6)P_3_ references were assigned accordingly based on concordant migration behavior and characteristic isotope mass shifts. Signals co-migrating within the [^18^O_6_]-Ins(1,3,4)P_3_ / Ins(1,4,5)P_3_ / Ins(1,4,6)P_3_ and [^18^O_6_]-Ins(1,3,5)P_3_ / Ins(2,4,6)P_3_ / Ins(2,3,5)P_3_ reference cluster were annotated conservatively, with Ins(1,4,5)P_3_ and Ins(2,3,5)P_3_ considered the most likely species based on relative abundance and established biosynthetic pathways. In cases where partial separation was achieved, peaks were annotated according to the corresponding reference species. Because the CE separation applied in this study is achiral, enantiomeric InsP_3_ pairs co-migrate and cannot be resolved; therefore, assigned signals represent either single isomers or unresolved enantiomeric pairs. All electropherograms were acquired under identical CE–ESI–MS conditions.

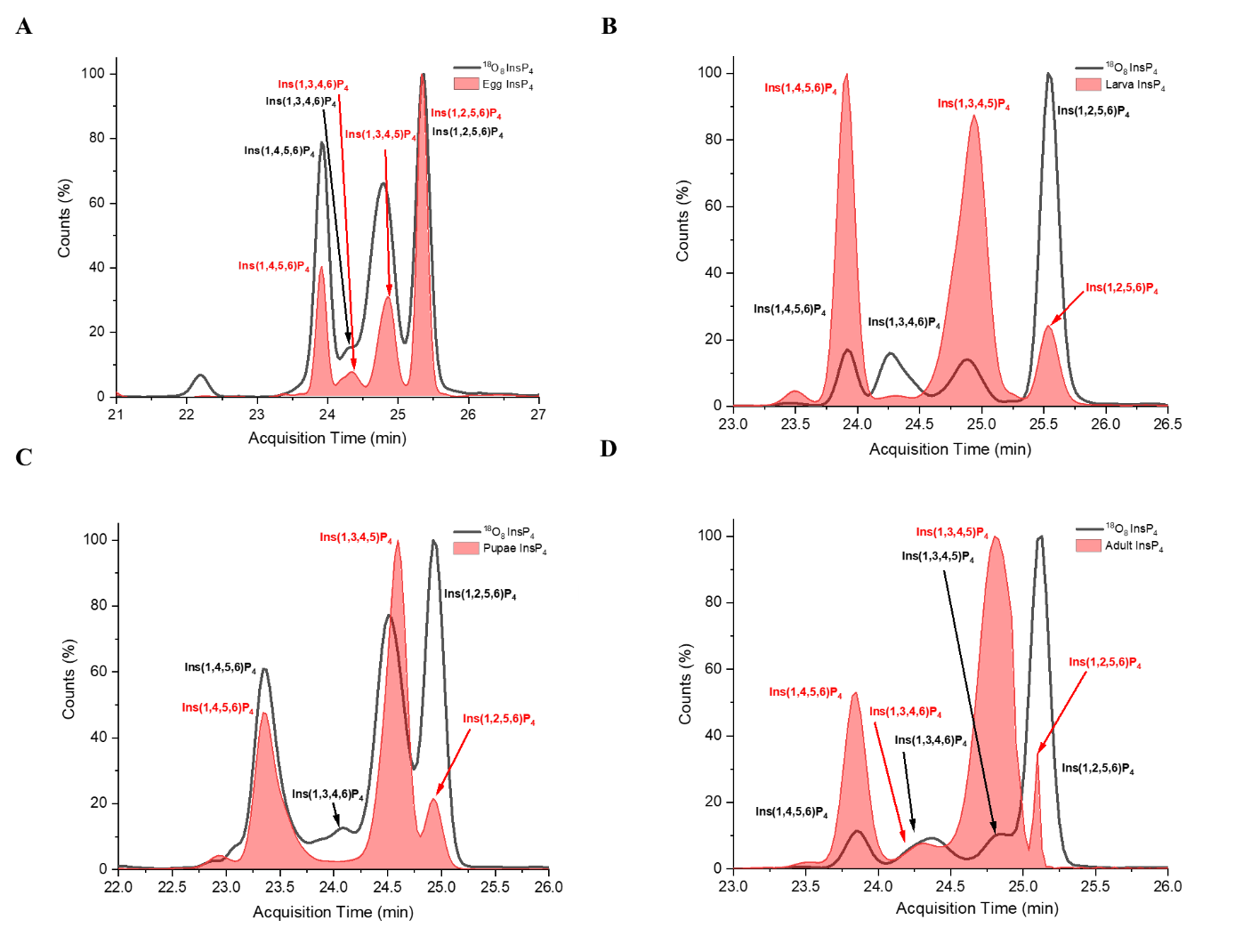

**Supplementary Figure S8. Assignment of InsP_4_ isomers in stage-specific *Drosophila melanogaster* samples using isotopically labelled references.** Extracted ion electropherograms showing positional assignment of InsP_4_ isomers in stage-specific *Drosophila* melanogaster samples. (**A**) egg, (**B**) larva, (**C**) pupae, and (**D**) adult extracts (red area) are overlaid with corresponding isotopically labeled reference standards analyzed under identical CE–ESI–MS conditions. InsP_4_ isomer assignment was achieved by comparing endogenous migration times with those of an [^18^O_8_]-InsP_4_ reference mixture, following prior calibration using [^13^C_6_]-InsP_4_ standards. Endogenous InsP_4_ species co-migrating with [^18^O_8_]-Ins(1,4,5,6)P_4_/[^18^O_8_]-Ins(3,4,5,6)P_4_, [^18^O_8_]-Ins(1,3,4,5)P_4_/[^18^O_8_]-Ins(1,3,5,6)P_4_, and [^18^O_8_]-Ins(2,3,5,6)P_4_/[^18^O_8_]-Ins(1,2,5,6)P_4_ references were assigned accordingly based on concordant migration behavior and characteristic isotope mass shifts. Additional InsP₄ signals migrating between reference peaks were conservatively annotated as consistent with Ins(1,3,4,5)P_4_ and/or enantiomers. Given the existence of 15 possible InsP_4_ positional isomers and the achiral nature of the CE separation, enantiomeric pairs cannot be resolved and alternative structural assignments cannot be excluded. All electropherograms were acquired under identical CE–ESI–MS conditions.

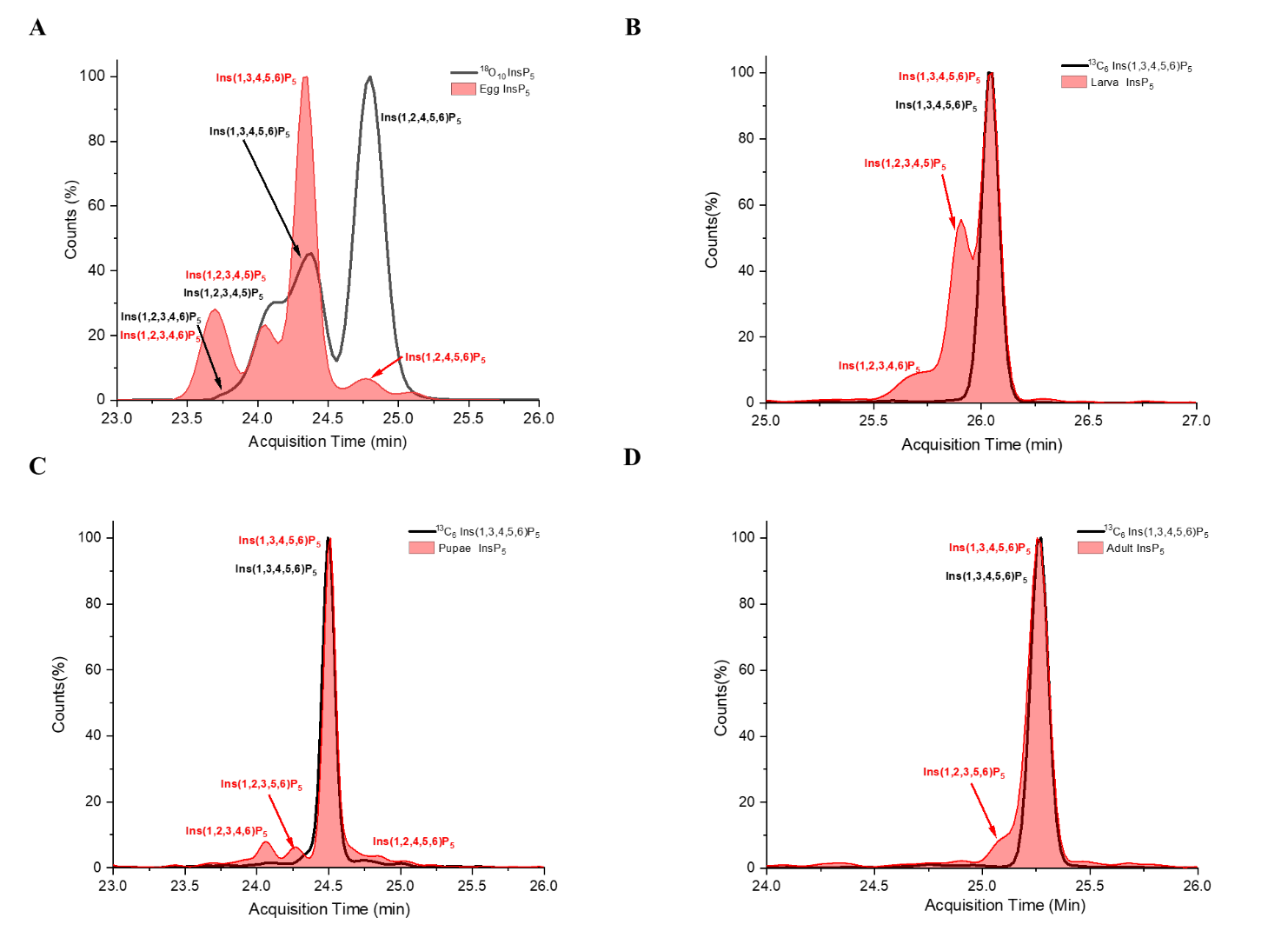

**Supplementary Figure S9. Assignment of InsP_5_ isomers in stage-specific *Drosophila melanogaster* samples using isotopically labelled references.** Extracted ion electropherograms showing positional assignment of InsP_5_ isomers in stage-specific *Drosophila* melanogaster samples. (**A**) egg, (**B**) larva, (**C**) pupae, and (**D**) adult extracts (red area) are overlaid with corresponding isotopically labelled reference standards analyzed under identical CE–ESI–MS conditions. InsP_5_ isomer assignment was achieved by comparing endogenous migration times with those of an [^18^O_10_]-InsP_5_ reference mixture, following prior calibration using [^13^C_6_]-InsP_5_ standards. Endogenous InsP_5_ species co-migrating with [^18^O_10_]‑Ins(1,2,3,4,6)P_5_, [^18^O_10_]-Ins(1,2,3,5,6)P_5_/[^18^O_10_]-Ins(1,2,3,4,5)P_5_, [^18^O_10_]-Ins(1,3,4,5,6)P_5_, and [^18^O_10_]-Ins(2,3,4,5,6)P_5_/[^18^O_10_]-Ins(1,2,4,5,6)P_5_ references were assigned accordingly based on concordant migration and characteristic isotope mass shifts. In panels (**B–D**), [^13^C_6_]-Ins(1,3,4,5,6)P_5_ (black trace) is overlaid with endogenous InsP_5_ signals (red area) to further confirm identity. Because the CE separation applied in this study is achiral, enantiomeric InsP_5_ pairs cannot be distinguished; therefore, reported signals represent either single isomers or unresolved enantiomeric pairs. All electropherograms were acquired under identical CE–ESI–MS conditions.

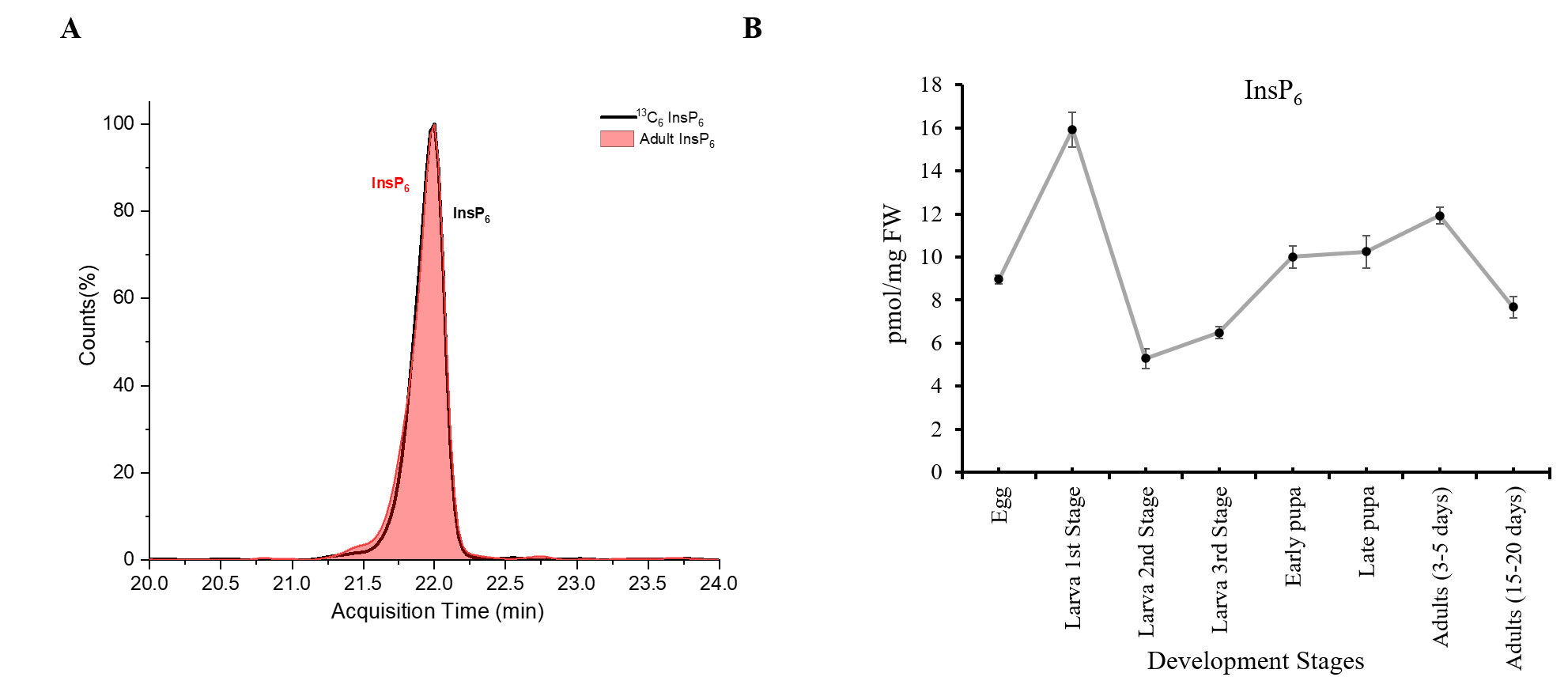

**Supplementary Figure S10. InsP_6_ detection and developmental dynamics in *Drosophila* melanogaster. (A)** Representative extracted ion electropherogram showing a co-migration of ^13^C_6_‑InsP_6_ (black trace) with endogenous *Drosophila* extracts InsP_6_ (red area) under CE–ESI–MS conditions.

**(B)** Quantitative analysis of InsP_6_ levels across developmental stages normalized to fresh tissue weight. InsP_6_ abundance exhibits stage-specific variation, with peak levels observed during larval development and reduced levels during pupation and adulthood (15-20 days). Data represent mean ± SD from three independent biological replicates.

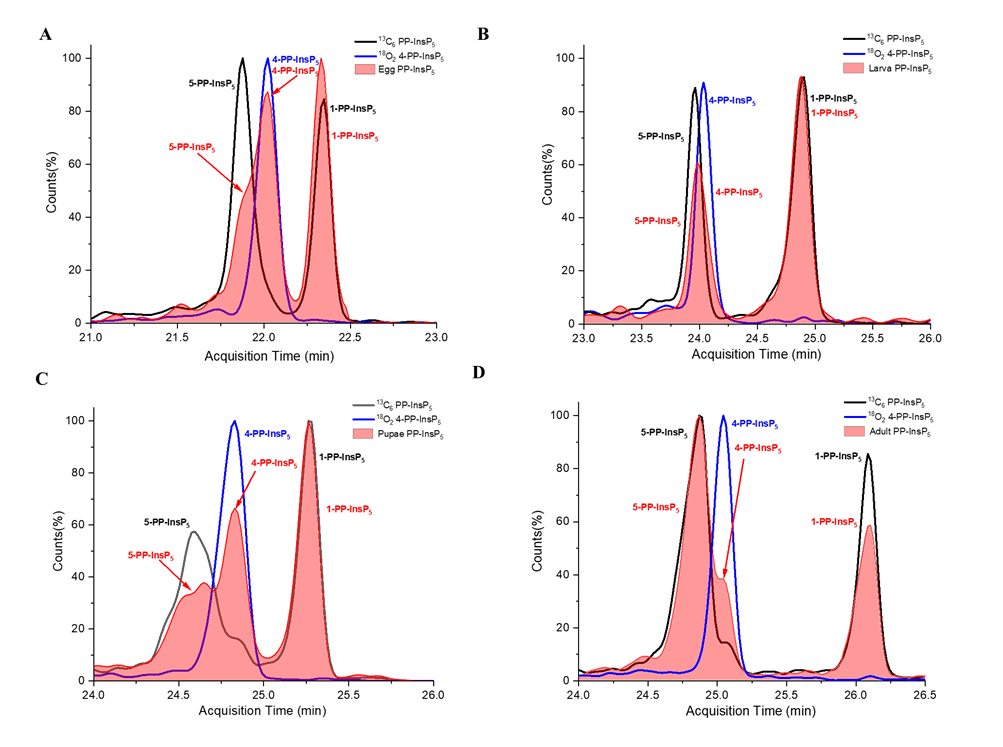

**Supplementary Figure S11. Assignment of PP-InsP_5_ isomers in stage-specific *Drosophila melanogaster* samples using isotopically labelled references.**

Extracted ion electropherograms showing positional assignment of PP-InsP_5_ isomers in stage-specific *Drosophila* *melanogaster* samples using isotopically labeled references. (**A**) egg, (**B**) larva, (**C**) pupae, and (**D**) adult extracts are shown (red area) overlaid with corresponding isotopically labeled PP-InsP_5_ standards (blue/black traces). Positional assignment was achieved by comparing migration times of endogenous PP-InsP_5_ signals with those of labeled reference standards analyzed under identical CE–ESI–MS conditions. Endogenous species co-migrating with labeled 5-PP-InsP_5_, 4-PP-InsP_5_ and 1-PP-InsP_5_, standards were assigned accordingly based on concordant migration behavior and characteristic isotope mass shifts. Because CE separation is achiral, enantiomeric diphosphoinositol phosphate species cannot be resolved; therefore, reported signals represent either single isomers or unresolved enantiomeric pairs. All electropherograms were acquired under identical CE–ESI–MS conditions.

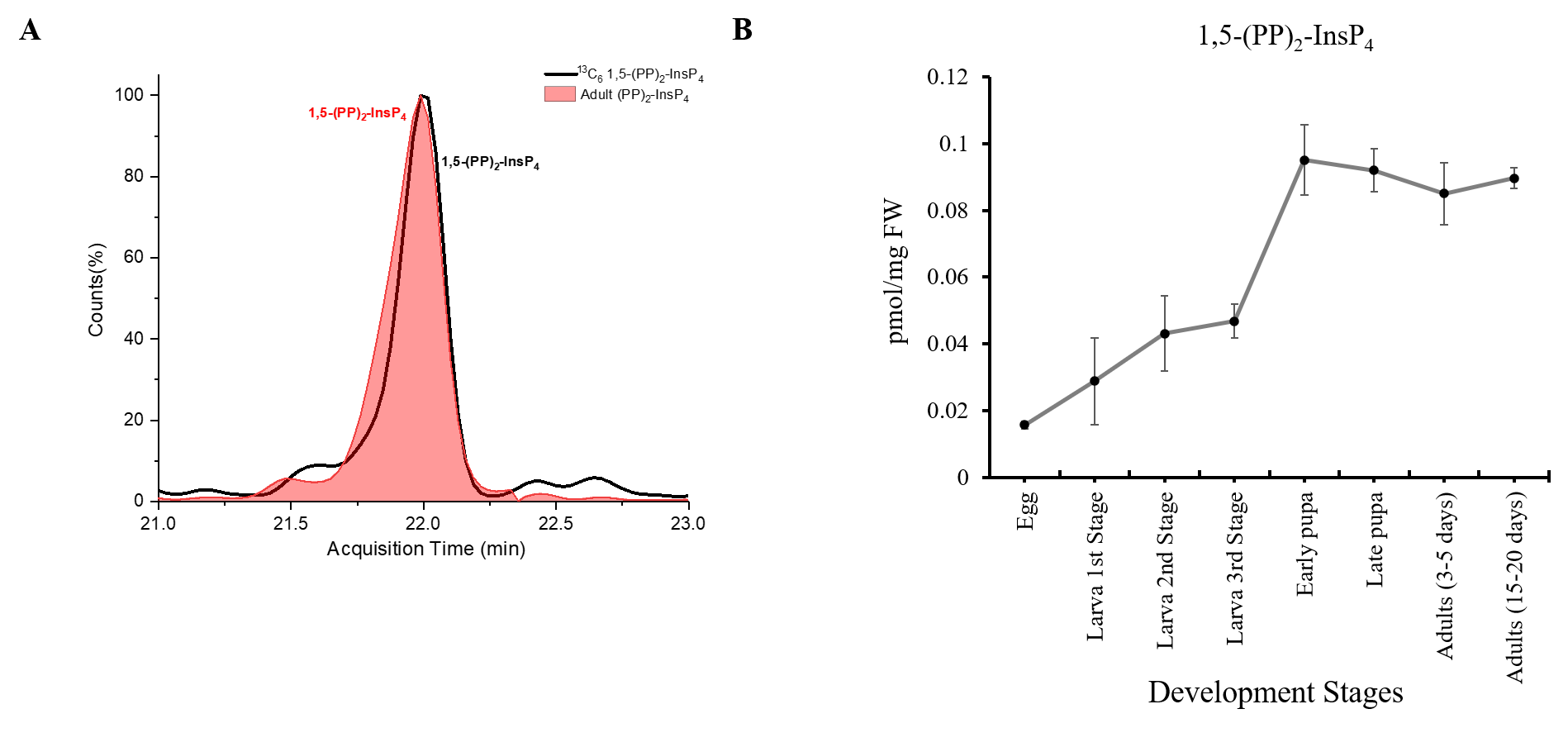

**Supplementary Figure S12. Developmental dynamics of 1,5-(PP)_2_-InsP_4_ in *Drosophila melanogaster*. (A)** Representative extracted ion electropherogram showing detection of 1,5-(PP)_2_-InsP_4_ in *Drosophila* *melanogaster* extracts (red area) under CE–ESI–MS conditions, co-migrating with ^13^C_6_ 1,5-(PP)_2_-InsP_4_ (black trace). **(B)** Quantitative analysis of 1,5-(PP)_2_-InsP_4_ across developmental stages normalized to fresh tissue weight. Levels increase during larval development and peak during early pupation, followed by stabilization in adults. Data represent mean ± SD from three independent biological replicates. Although 4,5-(PP)_2_-InsP_4_ may be present, its signal was below the quantitation limit under the applied analytical conditions and was therefore not included in quantitative measurements.

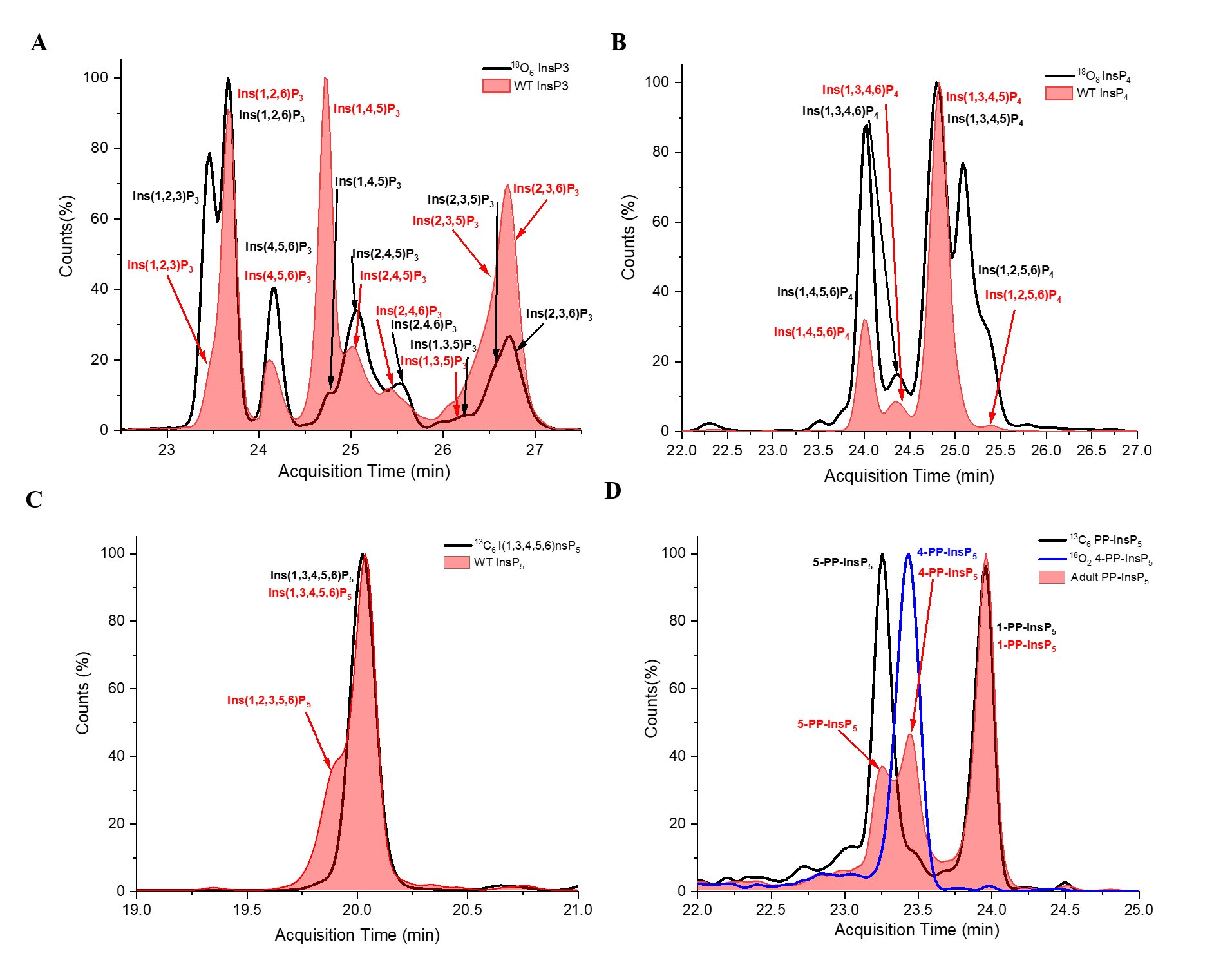

**Supplementary Figure S13. Representative electropherograms of InsP_3_ to PP-InsP_5_ species detected in WT Drosophila melanogaster.** Extracted ion electropherograms showing representative positional isomer profiles of (**A**) InsP_3_, (**B**) InsP_4_, (**C**) InsP_5_, and (**D**) PP-InsP_5_ species detected in WT *Drosophila melanogaster* pupae (12-24 h) extracts under identical CE–ESI–MS conditions. Annotated peaks indicate assigned positional isomers based on co-migration with isotopically labeled standards and validated pyro hydrolysis mixtures. InsP_3_ and InsP_4_ panels demonstrate extensive positional diversity across lower inositol phosphates. InsP_5_ is dominated by Ins(1,3,4,5,6)P_5_, with minor Ins(1,2,3,4,5)P_5_. The PP-InsP_5_ panel shows clear separation of diphosphoinositol phosphate isomers, including 5-PP-InsP_5_, 4-PP-InsP_5_ and 1-PP-InsP_5_. All signals represent extracted ion electropherograms acquired under identical analytical conditions. Because the CE separation is achiral, enantiomeric species cannot be resolved.

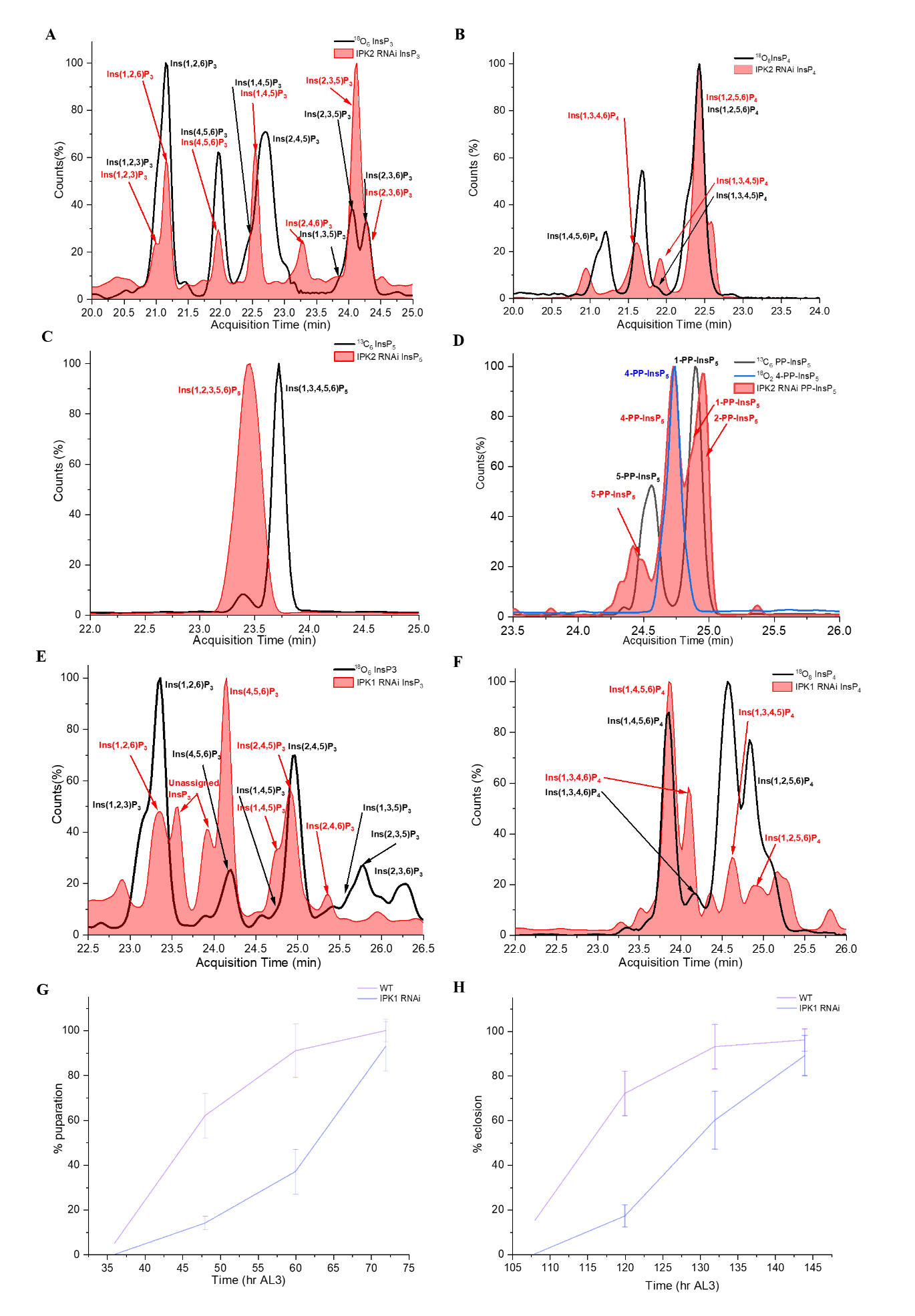

**Supplementary Figure S14. InsP positional isomers during IPK2 and IPK1 depletion and development timing for IPK1 depletion in *Drosophila melanogaster*.** Extracted ion electropherograms showing representative positional isomer distributions of IPK2 RNAi 12–24 h pupal extracts **(A)** InsP_3_, **(B)** InsP_4_, **(C)** InsP_5_, and **(D)** PP-InsP_5_ species (**E–F**) IPK1 RNAi 12–24 h pupal extracts under identical CE–ESI–MS conditions. Panels display profiles of InsP_3_, InsP_4_, and InsP_5_ species. Annotated peaks indicate assigned positional isomers based on co-migration with isotopically labeled standards and validated pyro hydrolysis mixtures. In IPK2-depleted and IPK1-depleted pupae, InsP_3_ and InsP_4_ panels demonstrate marked positional remodeling, with altered relative abundance of multiple low-abundance isomers alongside dominant species. InsP_4_ profiles show redistribution among Ins(1,3,4,5)P_4_, Ins(1,4,5,6)P_4_, Ins(1,2,5,6)P_4_, and related species. Notably, InsP_5_ profiles in IPK2 RNAi pupae are dominated by Ins(1,2,3,5,6)P_5_. For PP-InsP_5_ , a clear separation of 5-PP-InsP_5_ and 4-PP-InsP_5_ was observed in IPK2-depleted pupae. Additional signals that did not conclusively co-migrate with available reference standards are not assigned. All electropherograms represent extracted ion signals acquired under identical analytical conditions.

(**G-H**) Developmental timing analysis reveals delayed puparation and adult emergence upon IPK1 depletion. The percentage of animals undergoing pupal formation (**G**) and eclosion (**H**) was scored over time. Data represent n = 100 flies per genotype. (AL3 : After Larva 3^rd^ stage)

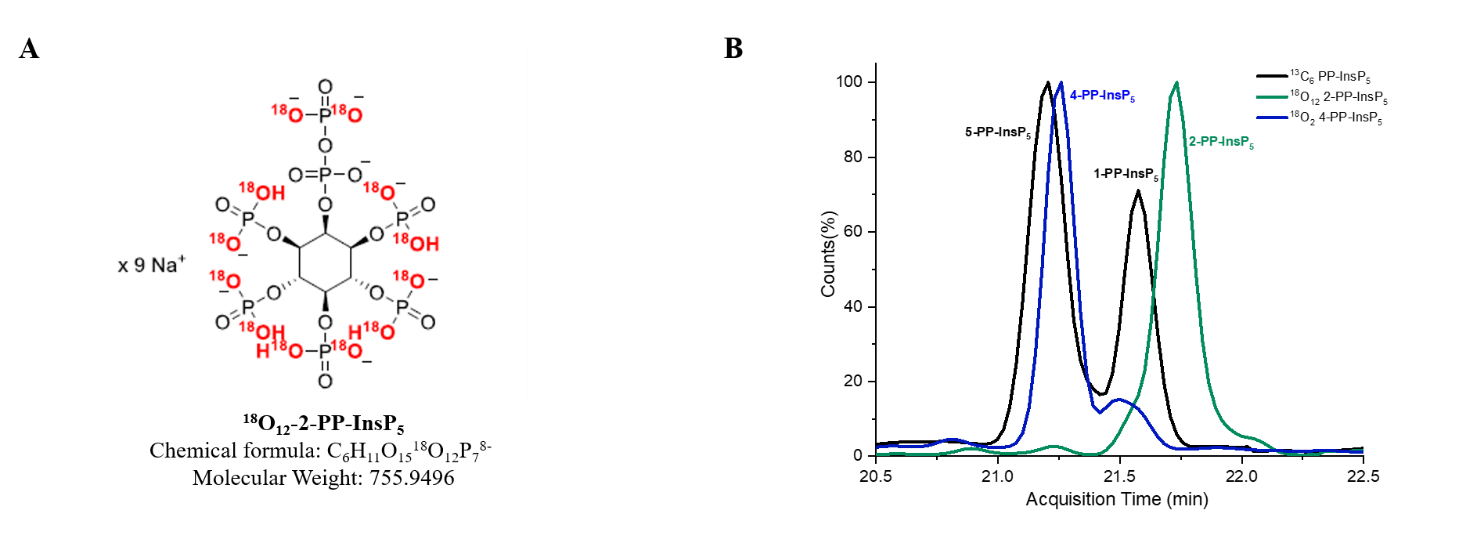

**Supplementary Figure S15. Synthesis and structural validation of ^18^O_12_-labeled 2-PP-InsP_5_ used for CE–MS assignment.**

(**A**) Structure of a newly synthesized [^18^O]_12_-2-PP-InsP_5_

(**B**) Representative CE-MS electropherograms showing separation of four PP-InsP_5_ isomers (5-PP-InsP_5_, 4-PP-InsP_5_, 1-PP-InsP_5_, and 2-PP-InsP_5_) under optimized CE conditions.

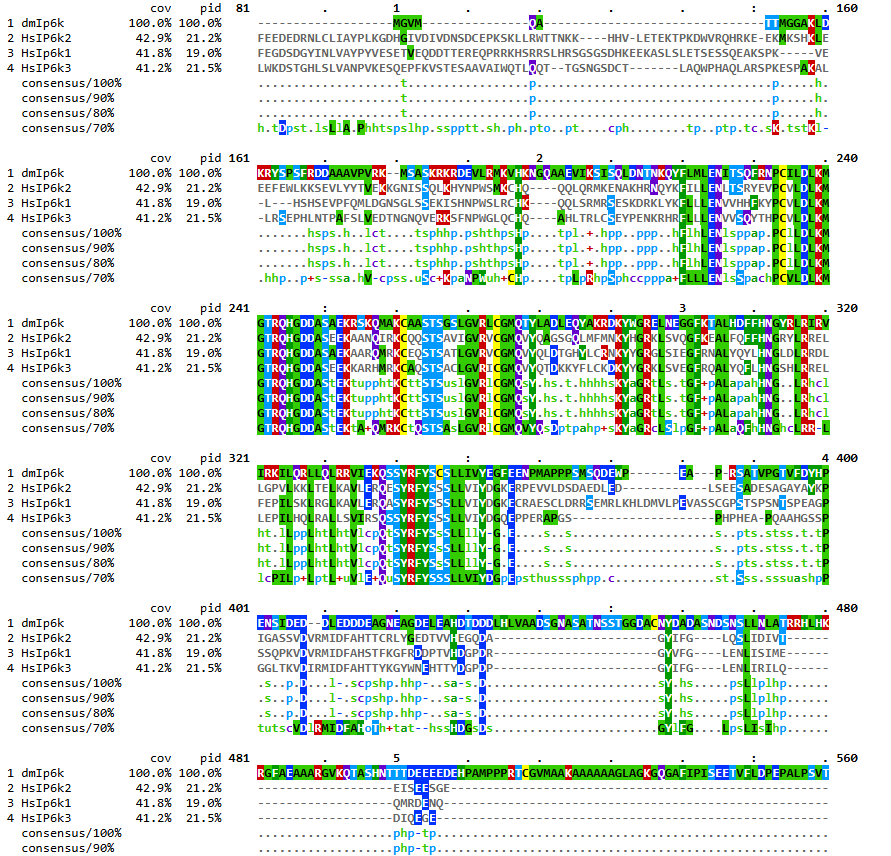

**Supplementary Figure S16. Sequence homology of IP6K proteins across species.**

Multiple sequence alignment of inositol hexakisphosphate kinase (IP6K) proteins from *Drosophila* *melanogaster* and human homologs (HsIP6K1, HsIP6K2, and HsIP6K3). Conserved residues and catalytic motifs characteristic of the IP6K kinase family are highlighted. The alignment reveals strong conservation within the catalytic core region, including the 233PxxxDxKxG241 motif and surrounding residues implicated in ATP binding and phosphate transfer, whereas the N- and C-terminal regions exhibit greater sequence divergence. Sequence identities are normalized to the aligned length.

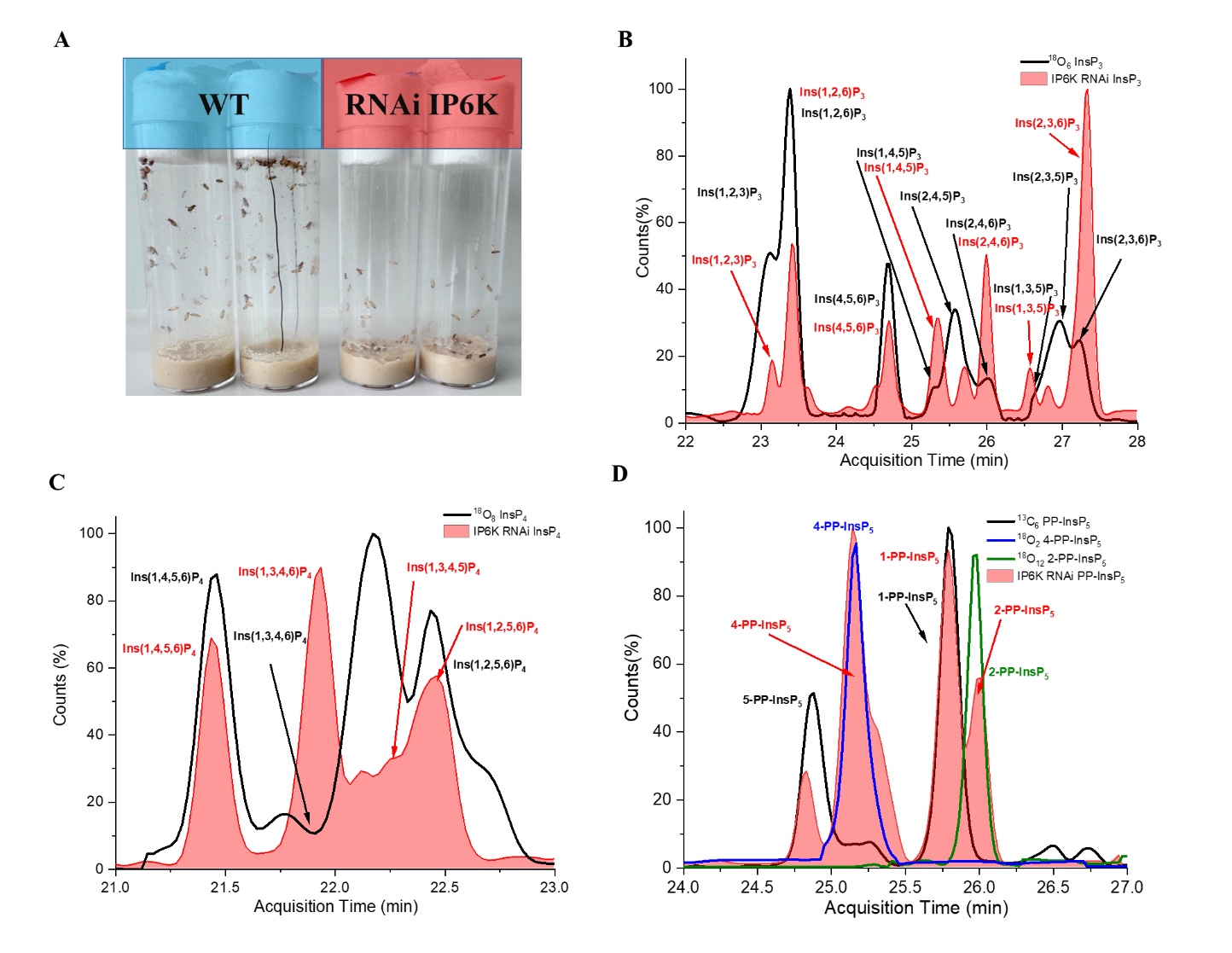

**Supplementary Figure S17. Phenotypic and metabolic consequences of IP6K depletion in *Drosophila melanogaster*. (A)** Representative images of wild-type (WT) and IP6K RNAi flies five days after eclosion. IP6K-depleted adults display impaired locomotion and frequently accumulate or collapse at the food surface. **(B)** Extracted ion electropherogram of InsP_3_ isomers detected in 12–24 h old IP6K RNAi pupae. Multiple positional InsP_3_ species are observed, indicating remodeling of lower inositol phosphates upon IP6K depletion. **(C)** Representative InsP_4_ electropherogram from 12–24 h old IP6K RNAi pupae, showing altered positional distribution compared to wild-type profiles. **(D)** PP-InsP_5_ electropherogram from 5-day-old IP6K RNAi adults. The expected 5-PP-InsP_5_ signal is markedly reduced, with redistribution toward alternative PP-InsP_5_ isomers, consistent with impaired IP6K-dependent phosphorylation activity. All electropherograms represent extracted ion signals acquired under identical CE–ESI–MS conditions.

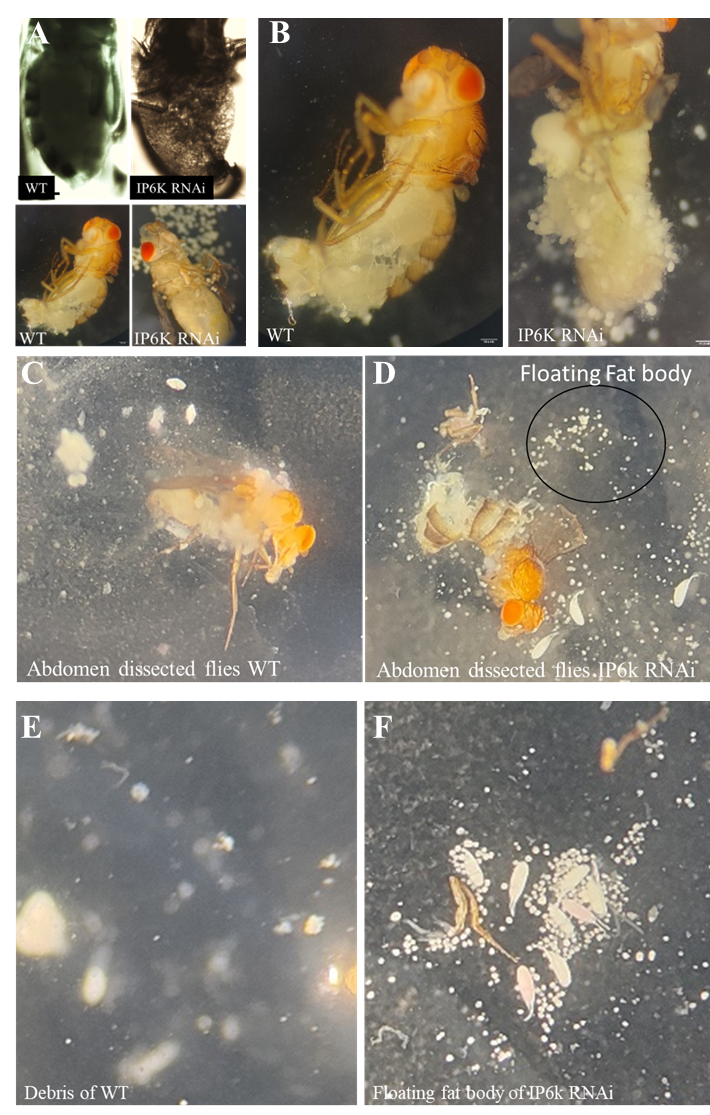
**Supplementary Figure S18. Persistence of floating larval fat body in IP6K-depleted adult *Drosophila melanogaster*.** (**A, B**) Representative abdomen images before and after dissection of adult and IP6K RNAi flies. **(C)** Wild-type (WT) adult abdomen showing normal tissue organization and absence of floating fat body cells. **(D)** IP6K RNAi adult abdomen displaying persistent floating fat body cells (circled). **(E)** Debris observed in WT dissections. **(F)** Enlarged view of floating fat body clusters in IP6K RNAi adults.

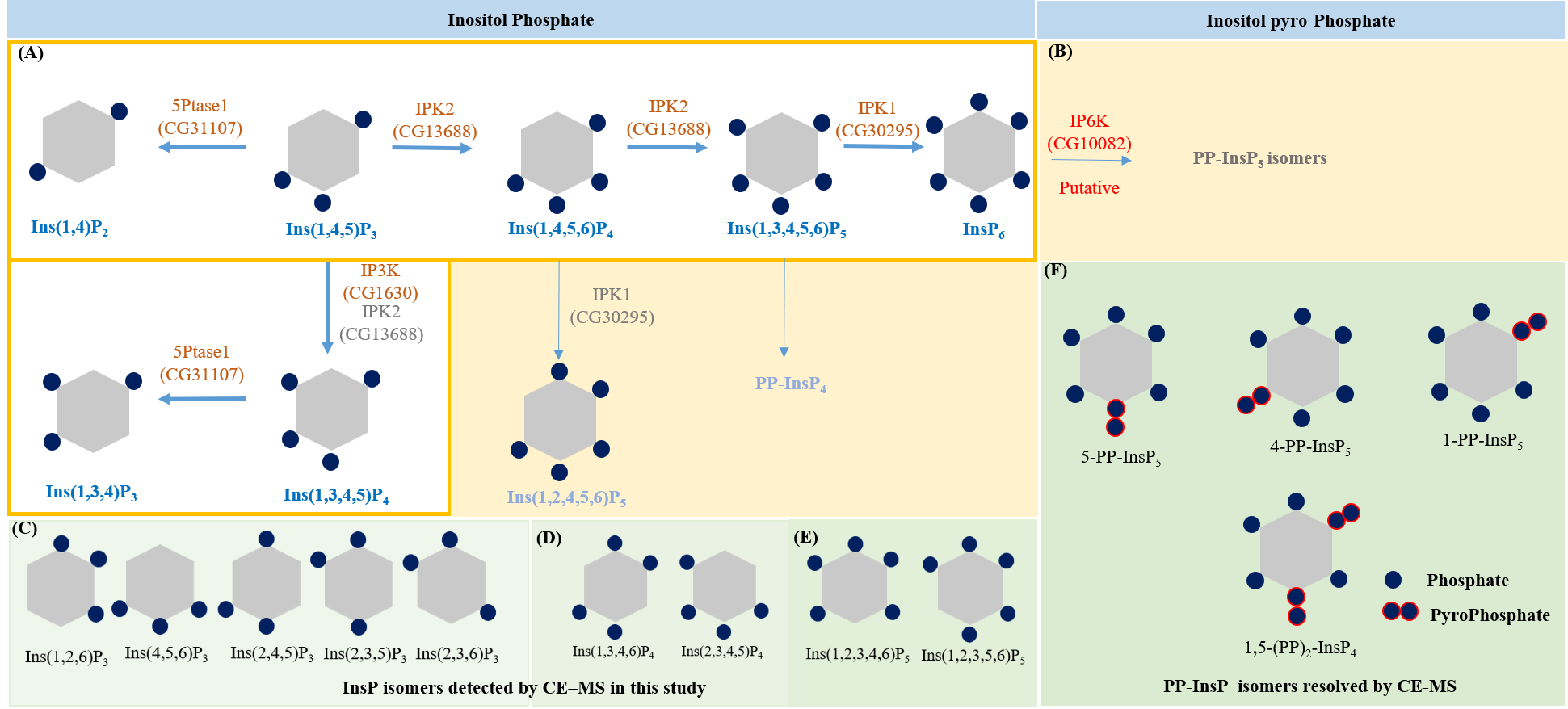

**Figure S19.** **Integrated map of inositol phosphate and pyrophosphate metabolism in *Drosophila melanogaster* incorporating CE-MS-resolved isomer diversity.**

Schematic representation of the inositol phosphate metabolic network illustrating the interconversion of InsP_2_ to InsP_6_ and their pyrophosphorylated derivatives. **(A**) is adapted from the genetic roadmap of inositol phosphate metabolism in *Drosophila* as proposed earlier^8^, describing the major route from Ins(1,4,5)P_3_ → Ins(1,4,5,6)P_4_ → Ins(1,3,4,5,6)P_5_ → InsP_6_ catalyzed by IPK2 (CG13688) and IPK1 (CG30295), with additional branches involving IP3K (CG1630) and inositol 5-phosphatase (CG31107). In this representation, solid arrows indicate metabolic steps directly supported by experimental evidence in the original publication^8^, whereas **(B**) represents the predicted part of the metabolic pathway proposed in ref.^8^. **(C-E**) summarize the inositol phosphate and pyrophosphate (PP-InsP) species detected in this study by CE–ESI–MS, including InsP_3_, InsP_4_, InsP_5_ and PP-InsP_5_ species. The lower panels illustrate the positional isomers of InsP_3_ to InsP_5_ resolved in this work, integrating the previously proposed pathway framework with the isomer-specific metabolite landscape identified in the present study, which is represented by a green color gradient.

**Supplementary Methods**

***Drosophila melanogaster* maintenance and developmental staging**

All experiments employed *Drosophila melanogaster*, with fly strains specified in Supplementary **Table S1**. Flies were kept in a standard cornmeal-yeast-agar medium (per 10 L: 74.5 g agar, 243 g dry yeast, 580 g corn flour, 552 ml molasses, 20.7 g Nipagin, 35 ml propionic acid) at 18–22 °C in a controlled laboratory setting.

For stage-specific developmental analyses, wild-type flies were used unless otherwise stated. All subsequent developmental analyses were performed at 22 °C. Adults were allowed to lay eggs on fresh medium for a defined time window, and eggs were collected within the first 0–6 h after egg laying (AEL) for developmental synchronization, after which adults were removed. Samples were subsequently collected at the following developmental windows: egg (0-4 h AEL), first-instar larvae (~24 h AEL), second-instar larvae (~48 h AEL), third-instar larvae (~72 h AEL), early pupae (0–12 h after pupariation), late pupae (48–72 h after pupariation), newly eclosed adults (3–5 days post-eclosion), and mature adults (15–20 days post-eclosion) without distinguishing between sexes. All samples were flash-frozen at the indicated stages and stored at −80 °C until extraction.

**Inositol Phosphates extraction and TiO₂ enrichment**

Inositol phosphates were extracted from 100–200 mg (wet weight) of flash-frozen whole *Drosophila melanogaster* samples collected at different developmental stages. All extraction steps were performed under ice-cold conditions. Samples were homogenized in 800–1,000 μL of chilled 1 M perchloric acid and incubated with gentle rotation at 4°C to precipitate proteins and release soluble metabolites. The homogenates were subsequently centrifuged at ≥16,000 × g for 15 min at 4°C. The resulting supernatants, containing soluble inositol phosphates, were transferred to fresh tubes.

Inositol phosphates and inositol pyrophosphates were enriched by TiO_2_ bead-based affinity purification. Briefly, 5 mg of TiO_2_ beads (Titansphere TiO_2_, 5 μm; GL Sciences) were washed sequentially with water and 1 M perchloric acid before being added to the extracts. Binding was allowed to proceed for 20 min at 4 °C with gentle rotation. Following incubation, the beads were collected by centrifugation at 5,000 × g and washed twice with ice-cold 1 M perchloric acid to remove non-specifically bound compounds.

Bound inositol phosphates were recovered by two sequential elutions with 10% ammonium hydroxide, each performed with gentle rotation for 5 min. The eluates were pooled and immediately dried under vacuum using a SpeedVac concentrator. The dried samples were stored at −20°C until reconstitution and analysis by capillary electrophoresis–mass spectrometry (CE–MS).

**Capillary Electrophoresis–Mass Spectrometry (CE–MS)**

**1. Chemicals and Reagents**

For our experiments, we used high-quality chemicals to ensure accuracy. Methanol, ammonium acetate, 2-propanol, and a 10% ammonia solution were all sourced from Carl Roth. The ^13^C_6_-labeled inositol phosphates and pyrophosphates we used were provided by Dorothea Fiedler at FMP Berlin. ^18^O-labeled inositol phosphates and pyrophosphates we used are made in house by chemical synthesis.

**2. Instrumentation and Setup**

All our analyses were done using an CE-ESI-QqQ system (Agilent 7100 CE with Agilent 6495C Triple Quadrupole and Agilent Jet Stream electrospray ionization source, adopting an Agilent CE-ESI-MS interface). For selected analyses, the CE system was alternatively coupled to a quadrupole time-of-flight (QToF) mass spectrometer using the same dedicated CE–MS adapter and sprayer kit. All measurements were performed in negative electrospray ionization mode. We used a bare fused-silica capillary (100 cm long, 50 µm internal diameter). Before starting, we flushed the capillary with 1 M NaOH for 10 minutes and then with water for another 10 minutes to prepare it.

Depending on targeted information, we used two different background electrolytes (BGEs):

BGE-1: 35 mM ammonium acetate, pH adjusted to 9.75 with ammonium hydroxide, for general profiling of inositol phosphates and pyrophosphates.

BGE-2: 40 mM ammonium acetate, pH adjusted to 9.08, for better separation of specific InsP_5_ and InsP_7_ isomers.

Before each run, we equilibrated the capillary with the chosen BGE for about 400 seconds. Samples (30 nL) were introduced by pressure injection (100 mbar for 15 seconds). For QToF measurements, we used a sheath liquid of water and 2-propanol (1:1, v/v) with continuous mass reference ions. For QqQ measurements, the sheath liquid was the same but without the reference ions. The sheath liquid flowed at 10 µL per minute. We controlled the instruments and collected data using Agilent’s Mass Hunter Workstation software (version 10.1). Details about the instrument settings can be found in **Table S4 and S5**.

**3. Quantitative analysis**

For quantitative analysis, dried TiO₂-enriched extracts were reconstituted in Milli-Q water prior to CE–MS measurement. Absolute and relative quantification of inositol phosphates was performed using stable isotope–labeled internal standards spiked directly into each sample before analysis. InsP_5_, InsP_6_, PP-InsP_5_, and (PP)_2_-InsP_4_ were quantified using known amounts of their corresponding ^13^C_6_-labeled standards as internal references. For lower inositol phosphates (InsP_3_ and InsP_4_), for which isotopically labeled standards are not commercially available for all positional isomers, quantification was performed using spiked ^13^C_6_- InsP_6_ as internal standard. Where available, positional ^13^C_6_-labeled Ins(1,2,3)P_3_ was additionally used to validate InsP_3_ quantification and enable direct comparison across lower InsP species under identical CE–MS conditions. This strategy ensured consistent normalization across the inositol phosphate series despite differences in available isotopic standards. Isotopic standards were spiked at concentrations specified in **Table S6**. These concentrations were selected to ensure accurate detection within the linear dynamic range of the instrument, enabling reliable and reproducible quantification.

**4. Standards, Assignment, and Quantification**

Confident positional assignment and quantification of inositol phosphate isomers were achieved using a complementary dual-isotope strategy involving ^13^C_6_- and ^18^O-labeled standards.

A reference library for lower InsPs positional isomers was generated by controlled heat dephosphorylation (pyrohydrolysis) of uniformly labeled ^18^O_12_-InsP_6_ (100 °C, 6 h) producing defined ^18^O-labeled InsP_3_–InsP_5_ species. The resulting ^18^O-labeled InsP_3_–InsP_5_ species retained identical electrophoretic migration behaviour compared to their unlabeled counterparts, while exhibiting predictable mass shifts that enabled unambiguous discrimination by high-resolution MS. Co-migration and isotope-specific mass signatures were used to validate peak identities.

Positional isomers were additionally validated by co-migration with ^13^C_6_-labeled inositol phosphate standards^9,10^. Following this validation, ^18^O-labeled reference compounds were used to assign inositol phosphate isomers in biological samples by matching both migration time and characteristic isotopic mass patterns.

For quantification, ^13^C_6_-Ins(1,2,3)P_3_ was included as an internal standard in all samples. Additional isotopically labeled standards were used as described in **Table S6**. Quantification was performed by comparing extracted ion electropherogram signal intensities relative to the internal standard under identical CE–MS conditions.

Because the capillary electrophoresis method applied in this study is achiral, enantiomeric resolution was not possible. Consequently, signals corresponding to enantiomeric pairs represent the combined abundance of both mirror-image species.

### 5. Analysis of inositol pyrophosphates

Inositol pyrophosphates were analyzed using the same CE–ESI–MS workflow. CE–MS resolved four PP-InsP_5_ positional isomers: 5-PP-InsP_5_, 1-PP-InsP_5_, 4-PP-InsP_5_ and 2-PP-InsP_5_ as well as a single detectable bis-diphosphorylated species, 1,5-(PP)_2_-InsP_4_. PP-InsP identities were confirmed by co-migration with available isotope-labeled reference standards and by the expected isotope mass shifts upon spiking^11,12^.

**6. MS validation and analytical robustness**

Fragmentation behaviour of selected inositol phosphate species was evaluated using targeted MS^2^ acquisition on the QToF instrument to confirm diagnostic fragmentation patterns. For sensitive and precise quantification, analyses were performed on a triple quadrupole (QqQ) mass spectrometer operating in multiple reaction monitoring (MRM) mode. The specific MRM transitions for each analyte are provided in **Table S5**.

To improve electrophoretic resolution of positional isomers, all samples were analyzed at least three times using two different background electrolytes specified above. This strategy enhanced separation across the InsP series and ensured robust detection and quantification of individual inositol phosphate species and their isomers.

**7. Data Analysis**

We extracted and integrated the peaks using MassHunter Quantitative Analysis software. The relative abundance of each compound was calculated directly from the integrated ion intensities. When comparing pathways, we expressed the results as percentages, using formulas:

Relative abundance of pathway A (%) = 100 × (Peak area of pathway A) / (Peak area of pathway A + Peak area of pathway B)

Relative abundance of pathway B (%) = 100 × (Peak area of pathway B) / (Peak area of pathway A + Peak area of pathway B)

**RNAi-Mediated Gene Silencing**

RNAi-mediated gene knockdown was performed using UAS-driven hairpin constructs obtained from the Vienna Drosophila Resource Center (VDRC). The following RNAi lines were used: IPK1 (CG30295) RNAi, KK-109497 (chromosome 2); IPK2/IPMK (CG13688) RNAi, GD-43825 (chromosome 2); and IP6K (CG10082) RNAi using two independent lines, KK-103749 and GD-38327 (both on chromosome 2).

RNAi induction was achieved using a heat-shock–inducible FLP/GAL4 system^13^. Briefly, crosses were established using the hsFlp[122]; tub-GAL4, UAS-mCD8::GFP driver line to enable inducible GAL4 activation following heat shock. This strategy allows temporal control of RNAi expression during development while maintaining consistent genetic background. GFP expression served as a marker for GAL4 activation.

The VDRC KK and GD RNAi libraries differ in design and potential off-target liabilities. KK lines employ short hairpin RNAs inserted at defined genomic loci and may, in rare cases, exhibit insertion-site-associated transcriptional artifacts. GD lines use longer dsRNA constructs inserted randomly in the genome, which may introduce low-level sequence-dependent off-target effects or incomplete knockdown efficiency. These factors were considered during interpretation of phenotypes.

To minimize potential RNAi artifacts, metabolic phenotypes were evaluated using high-resolution CE–MS profiling of individual inositol phosphate and diphosphoinositol phosphate isomers, providing pathway-specific biochemical signatures. In addition, CG10082 (IP6K) depletion was validated using two independent RNAi lines from different VDRC libraries, both of which produced consistent metabolic and developmental phenotypes.

### RNAi perturbations and sample selection for CE–MS analysis

Systemic RNAi perturbations targeting key inositol phosphate kinases were analyzed at defined developmental windows selected based on wild-type metabolic remodeling. For pupal analyses, individuals aged 12–24 h after pupation formation were collected. For adult analyses, flies aged 3–5 days post-eclosion were used. All RNAi samples were processed in parallel with matched controls using identical extraction, enrichment, and CE–MS workflows to enable direct comparison of InsP and PP-InsP profiles.

**Adult Dissection and Tissue Morphology Analysis**

For assessment of adult tissue organization, 3–5-day-old adults were anesthetized using CO₂ and dissected in ice-cold 1× phosphate-buffered saline (PBS). The abdominal cuticle was carefully opened using fine forceps under a stereomicroscope to expose internal tissues. Samples were examined immediately by bright-field microscopy. Particular attention was given to the organization of the abdominal adipose tissue. In control animals, the adult fat body formed organized lobular sheets attached to the body wall. In IP6K RNAi adults, tissue morphology was evaluated based on the presence of dispersed, unbound cellular clusters consistent with a disorganized or floating fat body–like phenotype. Representative images were acquired using identical exposure settings for control and experimental samples.

### Biological replicates and data reporting

For developmental profiling, each stage was analyzed using 3–5 independent biological replicates. RNAi perturbation experiments were performed using independently prepared biological replicates. Data are presented as mean values across biological replicates, with variability indicated as specified in the figure legends.
